# Turbulence orchestrates actin-mitochondria dynamics to preserve GPIbα and support iPSC-derived platelet biogenesis

**DOI:** 10.1101/2025.05.22.655481

**Authors:** Emiri Nakamura, Yasuo Harada, Sou Nakamura, Kosuke Fujio, Koenraad De Wispelaere, Chantal Thys, Kathleen Freson, Takuya Yamamoto, Akira Sawaguchi, Shiro Suetsugu, Koji Eto

## Abstract

The *in vitro* manufacturing of induced pluripotent stem cell-derived platelets (iPSC-platelets) remains limited by low yield and poor quality, thus impeding practical clinical application. To address this, we developed a turbulence-assisted system using defined physical stimulation to enhance platelet production from immortalized megakaryocyte progenitor cell lines (imMKCLs). This approach significantly increases the yield of glycoprotein Ibα (GPIbα)⁺ functional iPSC-platelets. However, the underlying mechanisms remained unclear. Here, we reveal that GPIbα⁺ iPSC-platelets retain high mitochondrial membrane potential, low phosphatidylserine (PS) exposure, and active ATP-dependent flippases, preserving mitochondrial integrity and platelet functionality. In contrast, GPIbα⁻ platelets exhibit PS exposure, ADAM17 activation, and selective GPIbα shedding. Notably, turbulent flow induces late-stage actin depolymerization, promoting even mitochondrial distribution within maturing imMKCLs and enhancing mitochondrial inheritance to iPSC-platelets. These findings highlight the key role of turbulence in regulating actin dynamics and mitochondrial allocation during platelet biogenesis, providing a mechanistic foundation for improving in vitro platelet production for therapeutic use.

## INTRODUCTION

Amid the scarcity of safe and immediately effective definitive therapies for patients with thrombocytopenia or acute bleeding, transfusion medicine using donor-derived platelet products remains the cornerstone of available treatments. However, these products face significant limitations, including a short shelf life of approximately 4–5 days (Brixner et al., 2024) due to the lack of long-term storage methods that simultaneously prevent the risk of bacterial growth and preserve platelet function and morphology, as well as an inconsistent supply that is entirely dependent on donors. A major clinical challenge is alloimmune platelet transfusion refractoriness (allo-PTR), which arises from alloimmunization due to previous transfusions or pregnancy (Panch et al., 2023). This condition is primarily driven by sensitization to platelet alloantigens, such as human leukocyte antigen class I (HLA-I) and human platelet antigen (HPA). Currently, there are no effective practical treatments when matched donor products for HLA-I or HPA are unavailable.

In this context, we conducted the first-in-human clinical trial of induced pluripotent stem cell-derived platelets (iPSC-platelets) as an autologous transfusion to an allo-PTR patient with anti-HPA-1a antibodies (Sugimoto et al., 2022a). The trial was made possible by developing immortalized megakaryocyte progenitor cell lines (imMKCLs) capable of generating iPSC-platelets (Nakamura et al., 2014). Following an established protocol for imMKCL generation, we successfully created a master cell bank of imMKCLs, enabling on-demand platelet production (Sugimoto et al., 2022b).

Another key breakthrough in platelet biomanufacturing was the development of an efficient, large-scale production system. A pivotal innovation in this field involved identifying a novel mechanobiological concept: turbulence synergizes with shear stress to promote megakaryocyte maturation and functional platelet biogenesis. This was inspired by *in vivo* observations where physical stress—generated by blood flow—enhanced platelet formation, a phenomenon subsequently recapitulated *in vitro*.

*In vivo*, multiple mechanisms of thrombopoiesis have been proposed, including proplatelet extension, cytoplasmic rupture, fragmentation, and membrane budding at the bone marrow sinusoidal interface (Nishimura et al., 2015; Tilburg et al., 2022; Potts et al., 2020). Moreover, recent discoveries have identified the pulmonary vasculature, especially at vessel bifurcations, as a significant site of platelet generation (Lefrançais et al., 2017; Zhao et al., 2023). These bifurcations induce turbulent flow, which can be mimicked in vitro using platforms like the VerMES reactor or even in simple flask cultures. Under such conditions, the number of GPIbα⁺ platelets markedly increases (Ito et al., 2018; Okamoto et al., 2024). However, the precise molecular mechanisms by which turbulence enhances GPIbα⁺ platelet production remain poorly understood, thereby preventing full optimization of *in vitro* systems.

While crucial for thrombosis, the N-terminal domain of GPIbα—the platelet receptor for von Willebrand factor (vWF) and fibrinogen—is also known to be shed by ADAM17. Despite this, the GPIb complex, comprising GPIbα, GPIbβ, GPV, and GPIX, remains structurally stable under shear stress caused by blood flow (Nishikii et al., 2008). In addition to ectodomain shedding, desialylation of the N-terminal region of GPIbα also leads to rapid clearance of platelets *in vivo* (Jansen et al., 2012). Therefore, wholly preserving the structural integrity of GPIbα is essential for maintaining the functional quality of platelets, even after *in vitro* production.

Mitochondria are essential for platelet function, primarily by generating ATP to support platelet activation and aggregation (Melchinger et al., 2019). They also regulate reactive oxygen species (ROS) levels and calcium homeostasis—both of which are critical for platelet activity (Chen et al., 2013; Kholmukhamedov et al., 2018; Ajanel et al., 2023). Mitochondrial dysfunction impairs platelet responses (Richman et al., 2023; Ding et al., 2023) and has been associated with various pathological conditions, including Parkinson’s disease, Alzheimer’s disease, and type 2 diabetes (Shi et al., 2008; Avila et al., 2012; Zharikov and Shiva, 2013; Antony et al., 2015; Wang et al., 2017; Twomey et al., 2018; Melchinger et al., 2019). These findings underscore the critical role of mitochondria in maintaining normal platelet function. However, our understanding of mitochondrial dynamics during platelet production remains limited. In particular, how mitochondria are evenly distributed to platelets during their biogenesis has not been investigated.

In this study, we revealed that functional platelet production was significantly increased under turbulent flow by enhancing balanced mitochondrial segregation. Specifically, GPIbα⁺ iPSC-platelets generated under turbulent conditions show elevated mitochondrial membrane potential and reduced phosphatidylserine (PS) exposure, indicating preserved mitochondrial function. In contrast, GPIbα⁻ iPSC-platelets, lacking sufficient ATP to fuel flippases, display increased PS exposure and undergo ADAM17-dependent GPIbα shedding. Furthermore, turbulence promotes balanced F-actin dynamics in megakaryocytes, supporting even mitochondrial distribution and facilitating efficient platelet biogenesis. These findings underscore the critical role of turbulence in orchestrating intracellular cytoskeletal remodeling and mitochondrial integrity, offering key insights into improving both the quality and yield of *in vitro* platelet production systems.

## RESULTS

### Turbulence enhances high-quality iPSC-platelet production indicated by increasing GPIbα expression and mitochondrial membrane potential

To determine whether varying levels of turbulent energy and shear stress influence GPIbα expression, we compared iPSC-platelet production from Clone 17 imMKCLs in different conditions: static culture (dish), orbital shaking flask (sub-optimal shear and turbulence), and the VerMES™ bioreactor (optimal shear and turbulence) (Ito et al., 2018) (Fig. 1 A; Fig. S1 A). By analyzing surface markers such as integrin alpha 2b (CD41) and GPIbα, we found that turbulence not only increases the functional iPSC-platelet yield but also improves the proportion of functional iPSC-platelets (Fig. 1, B–D). Only GPIbα⁺ iPSC-platelets were activated by ADP/Trap6 or phorbol 12-myristate 13-acetate (PMA) stimulation (Fig. S1 B) and fluorescent upon Calcein-AM incubation, indicating high viability (Fig. 1 E). Interestingly, GPIbα⁺ iPSC-platelets showed higher mitochondrial membrane potential (MMP), as revealed by MitoTracker Deep Red staining, compared to GPIbα⁻ iPSC-platelets (Fig. 1 F). Additionally, a culture swap experiment (dish to flask, flask to dish on day 4) revealed that turbulence from day 4 onward, aligned with the onset of iPSC-platelets production (Fig. S1, E and F), is essential for improved iPSC-platelet yield (Fig. 1, G and H), suggesting a positive relationship between mitochondrial health and GPIbα expression that is promoted by turbulence after day 4.

**Figure 1.**
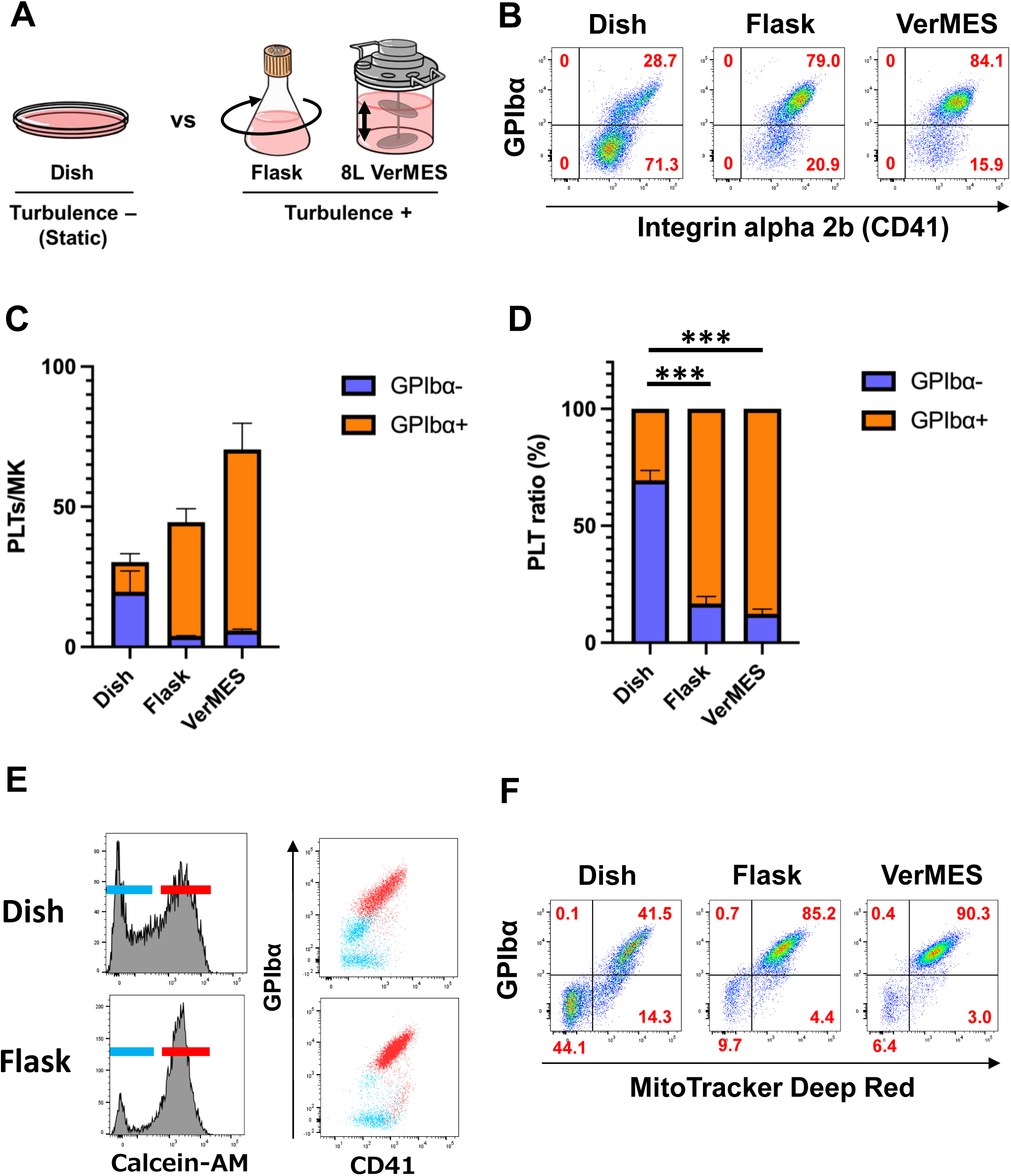
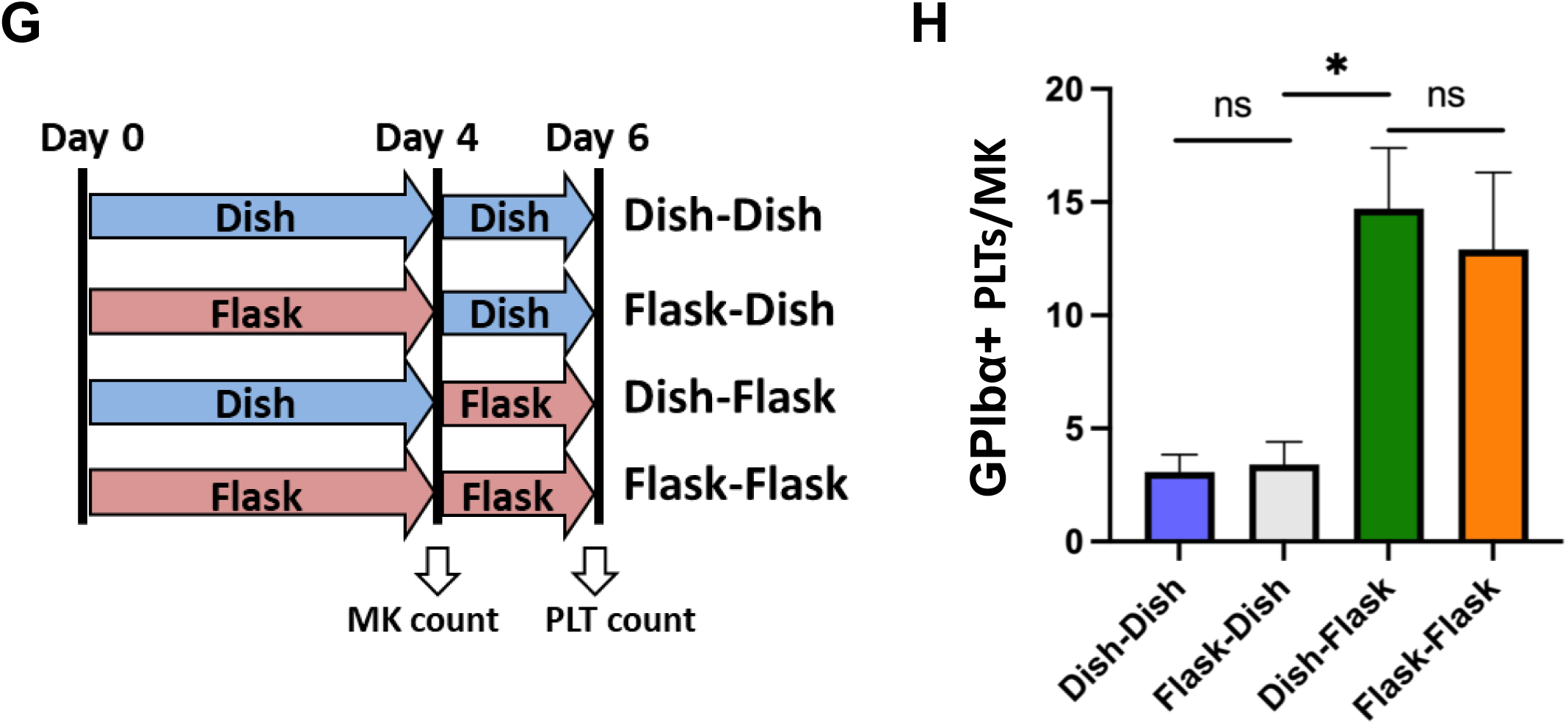
**Turbulence enhances high-quality iPSC-PLT production, as indicated by increased GPIbα expression and mitochondrial membrane potential** (A) Schematic illustration of three culture systems: dish (static), flask (suboptimal turbulence), and VerMES (optimal turbulence). (B–D) Analysis of integrin αIIb (CD41) and GPIbα expression on day 6 platelets derived from each culture condition. Platelets were defined as CD41⁺ particles. (B) Representative flow cytometry plots of platelets generated under dish (left), flask (middle), and VerMES (right) conditions. (C) Quantification of the number of GPIbα⁺ (orange) and GPIbα⁻ (blue) platelets produced per megakaryocyte. (D) Quantification of the percentage of GPIbα⁺ (orange) and GPIbα⁻ (blue) platelets. Data are presented as mean ± SEM (n = 3). Statistical significance was determined using an unpaired two-tailed t-test (***p < 0.001). (E) Representative flow cytometry plots showing Calcein-AM fluorescence (left) and surface expression of GPIbα and CD41 (right) in day 6 platelets cultured in a dish (top) or flask (bottom). (F) Representative flow cytometry plots showing mitochondrial membrane potential (MitoTracker Deep Red) and GPIbα expression in day 6 platelets cultured in a dish (left), flask (middle), or VerMES (right). (G, H) Platelet production per megakaryocyte in a day 4 culture swap experiment. (G) Schematic diagram of the experimental design for culture condition swapping. (H) Quantification of platelets produced per megakaryocyte under the following conditions: dish-dish, flask-dish, dish-flask, and flask-flask. Data are presented as mean ± SEM (n = 3). Statistical analysis was performed using an unpaired two-tailed t-test (ns, not significant; *p < 0.05).

### Mitochondria activity regulates GPIbα expression

Next, we investigated whether mitochondria play a positive regulatory role in GPIbα expression. We compared MMP, reactive oxygen species (ROS) levels, and ATP levels in GPIbα⁺ versus GPIbα⁻ iPSC-platelets and found that GPIbα⁺ iPSC-platelets exhibited higher levels across all three parameters (Fig. 2 A). Furthermore, MMP was inversely correlated with phosphatidylserine (PS) exposure, measured by annexin V binding, a marker of apoptosis (Fig. 2 B). These findings were consistent in the produced iPSC-platelets from imMKCLs transfected with CoralHue mito-mAG vector, which allowed visualization of the reduction in intact mitochondria in static iPSC-platelets compared to those cultured in flasks (Fig. 2 C; Fig. S2 A).

**Figure 2.**
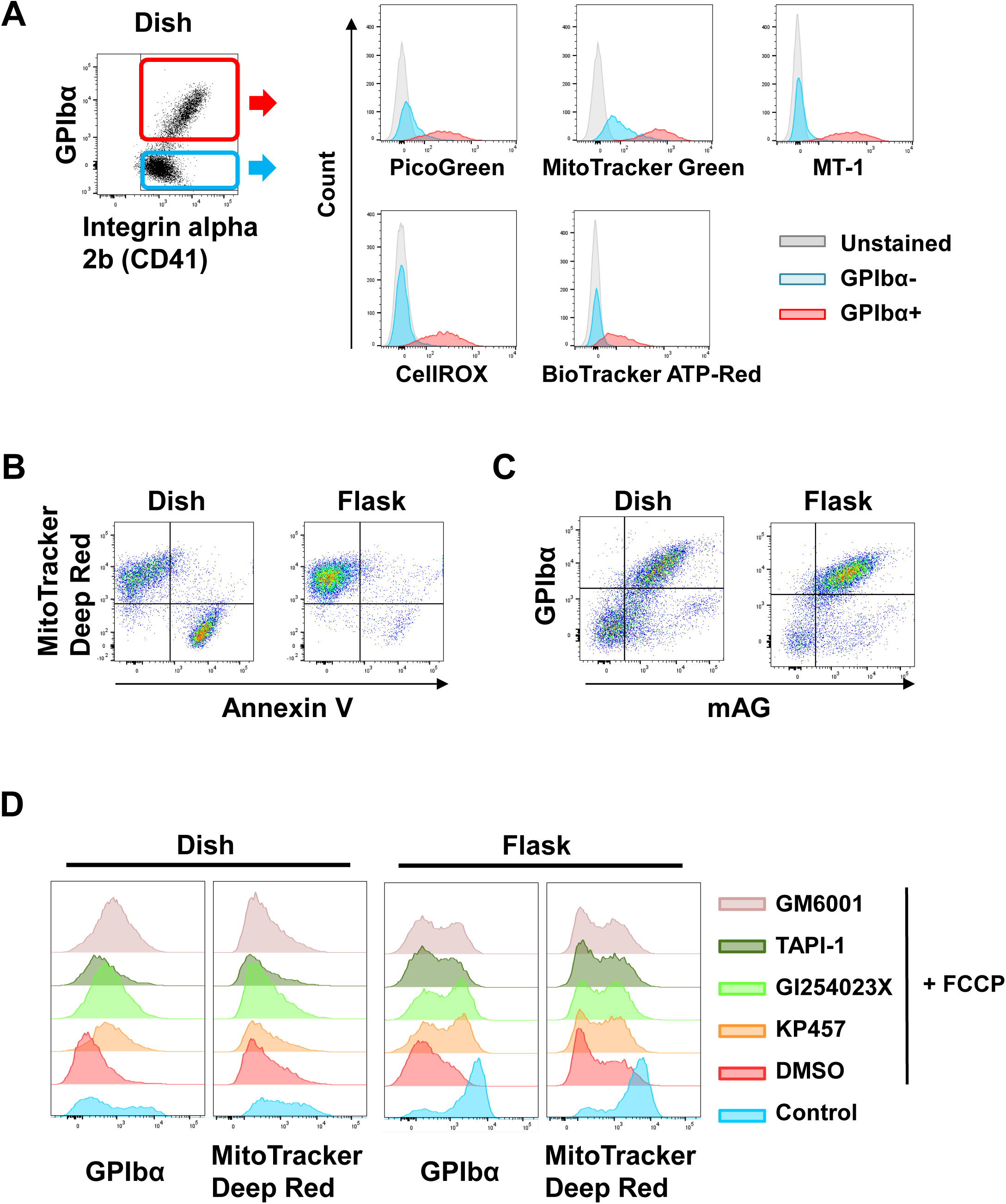
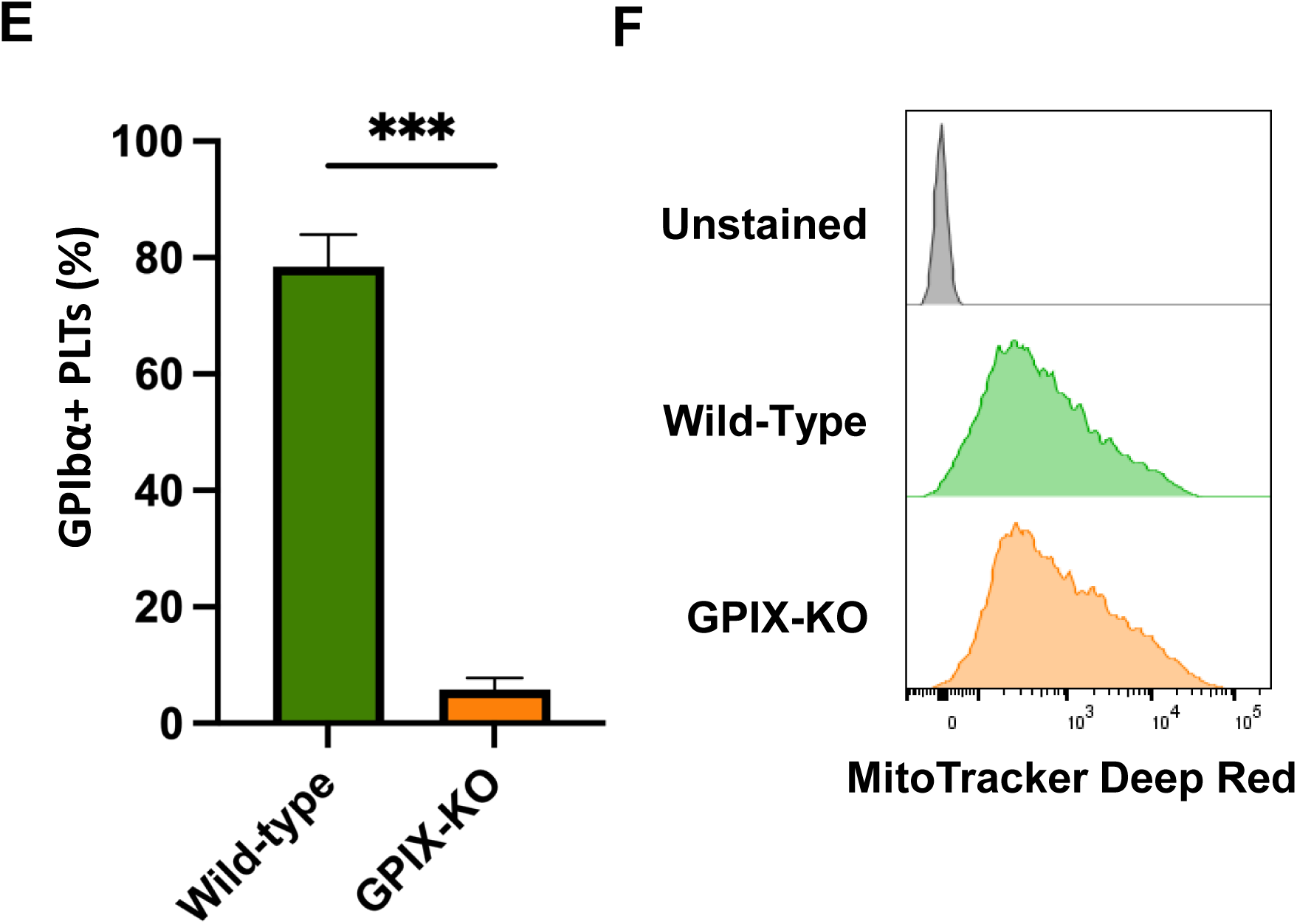
**Mitochondrial activity regulates GPIbα expression** (A) Representative flow cytometry plots and histogram overlays showing staining for mitochondrial DNA (PicoGreen), mitochondrial content (MitoTracker Green), mitochondrial membrane potential (MT-1), reactive oxygen species (ROS; CellROX), and adenosine triphosphate (ATP; BioTracker ATP Red) in GPIbα⁺ (red) and GPIbα⁻ (blue) day 6 platelets cultured under static conditions. Unstained controls are shown in gray. (B) Representative flow cytometry plots showing mitochondrial membrane potential (MitoTracker Deep Red) and phosphatidylserine exposure (annexin V) in day 6 platelets cultured in a dish (left) or flask (right). (C) Representative flow cytometry plots showing GPIbα staining and mitochondria-specific mAzami Green fluorescence in day 6 platelets derived from dish (left) or flask (right) cultures. (D) Representative histograms showing GPIbα expression (far left, middle right) and mitochondrial membrane potential (MitoTracker Deep Red; middle left, far right) in control (blue) and after treatment with DMSO (red), KP457 (orange), GI254023X (green), TAPI-1 (dark green), or GM6001 (purple) in day 6 platelets prior to FCCP treatment, from dish (left) or flask (right) cultures. (E, F) Comparison of platelet production and mitochondrial membrane potential between wild-type and GPIX-KO imMKCLs. (E) Quantification of the percentage of CD41⁺/GPIbα⁺ platelets produced by wild-type (green) or GPIX-KO (orange) imMKCLs. Data are presented as mean ± SEM (n = 3). Statistical significance was determined using an unpaired two-tailed t-test (***p < 0.001). (F) Representative histograms showing mitochondrial membrane potential in platelets derived from wild-type (green, middle) or GPIX-KO (orange, bottom) imMKCLs. Unstained control is shown in gray (top).

To further investigate the role of mitochondria in GPIbα expression, we inhibited mitochondrial activity using FCCP (carbonyl cyanide-p-trifluoromethoxyphenylhydrazone), a well-known uncoupler of proton-mediated ATP production in mitochondria. This treatment led to a decrease in GPIbα expression, which was reversed when iPSC-platelets were pre-treated with ADAM17 inhibitors such as KP457 (Hirata et al., 2017), high concentration of GI254023X (Metz et al., 2012), TAPI-1 (Nishikii et al., 2008), and GM6001 (Bergmeier et al., 2004) (Fig. 2 D). These results were also observed in donor-derived human platelets (Fig. S2 F).

Additionally, to confirm whether MMP levels are dependent on the GPIb-X-IX complex, we used GPIX-knockout (KO) imMKCLs, which predominantly produce GPIbα⁻ iPSC-platelets. However, no significant difference in MMP was observed between these and control cells (Fig. 2, E and F), thus confirming that mitochondria are a positive upstream regulator of GPIbα.

### Mitochondria supply flippases with ATP to maintain GPIbα expression

It has been previously reported that flippases mediate PS exposure, which in turn activates ADAM17 (Sommer et al., 2016), and that scramblases, such as TMEM16F (Fujii et al., 2015), become active during platelet functional decline. To determine whether mitochondrial regulation of GPIbα expression involves these factors, we examined PS and phosphatidylcholine (PC) dynamics in iPSC-platelets using fluorescent PS (NBD-PS) and PC (NBD-PC), which are internalized by recipient platelets when flippases or scramblases are active, respectively (Fig. 3 A). Consistent with our earlier findings using the annexin V binding assay (Fig. S1 C), NBD-PS was internalized by GPIbα⁺ but not GPIbα⁻ iPSC-platelets (Fig. 3 B; Fig. S3 A). When ionomycin was added, a significant increase in calcium levels and a subsequent decrease in ATP levels were observed, resulting in reduced GPIbα expression and diminished NBD-PS internalization (Fig. 3, C and D; Fig. S3 A), indicating that mitochondrial ATP production fuels flippases, internalizing PS and maintaining the inactive conformation of ADAM17, preventing GPIba shedding in iPSC-platelets.

**Figure 3.**
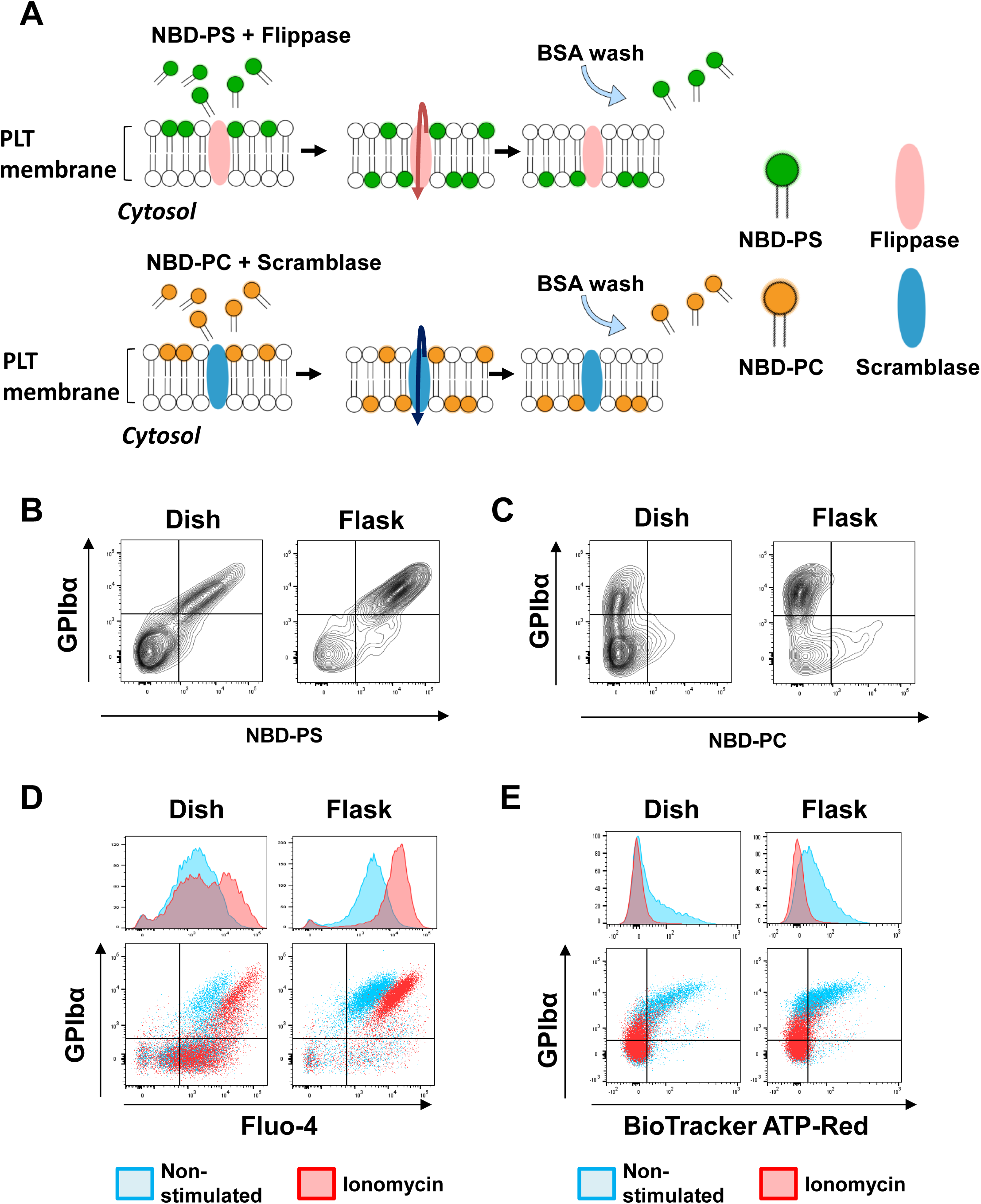
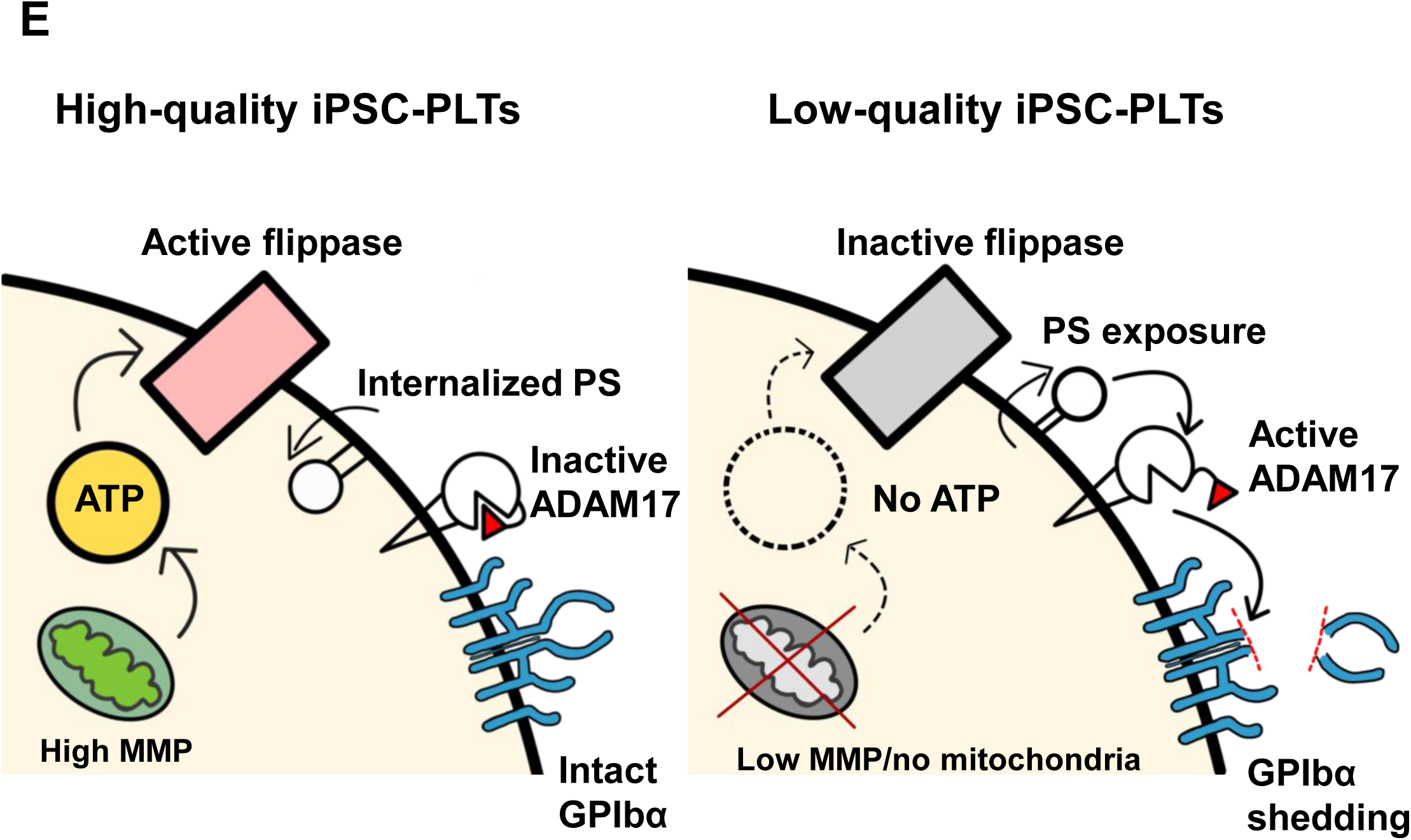
**Mitochondrial ATP production supports flippase-mediated membrane asymmetry** (A) Schematic illustration of the assay used to evaluate flippase and scramblase activity. (B, C) Representative flow cytometry plots showing surface GPIbα expression and NBD-labeled phospholipid incorporation in day 6 platelets cultured in a dish (left) or flask (right). (B) NBD-phosphatidylserine (NBD-PS); (C) NBD-phosphatidylcholine (NBD-PC). (D, E) Flow cytometric analysis of GPIbα expression, intracellular calcium levels, and ATP content in day 6 platelets derived from dish and flask cultures. (D) Representative plots and histograms showing surface GPIbα expression and intracellular calcium levels (Fluo-4) in platelets from dish (left) and flask (right) conditions. (E) Representative flow cytometry plots showing surface GPIbα expression and intracellular ATP levels (BioTracker ATP Red) in platelets cultured in a dish (left) or flask (right). (F) Schematic model illustrating differences in GPIbα expression between high-quality (left) and low-quality (right) iPSC-platelets.

### Turbulence imMKCLs exhibit uniform mitochondria distribution

Next, we investigated intracellular differences between maturing dish and flask imMKCLs, as mouse and human platelets inherit their mitochondria from megakaryocytes (Richardson et al., 2005; Jacob et al., 2024). After staining mitochondria with MitoTracker Deep Red and F-actin with SPY555-FastAct, a F-actin-stabilizing jasplakinolide derivative, respectively, while confocal imaging revealed uniform distribution of mitochondria and F-actin in flask imMKCLs, dish imMKCLs displayed distinct megakaryocyte polarization. However, no differences were observed in the surface expression of CD41/GPIbα or mitochondrial membrane potential (MMP) over the 6-day culture period (Fig. 4, A and B; Fig. S4, A and B). When mitochondrial distribution was disrupted using Mdivi-1, there was a dose-dependent shift toward GPIbα⁻ iPSC-platelet production (Fig. 4 C). Although no significant differences were observed in the oxygen consumption rate between day 4 dish or flask imMKCLs, differences in MMP were only seen in iPSC-platelets (Fig. 4, D and E), suggesting that turbulence promotes uniform mitochondrial distribution in megakaryocytes leading to efficient mitochondria inclusion by platelets.

**Figure 4.**
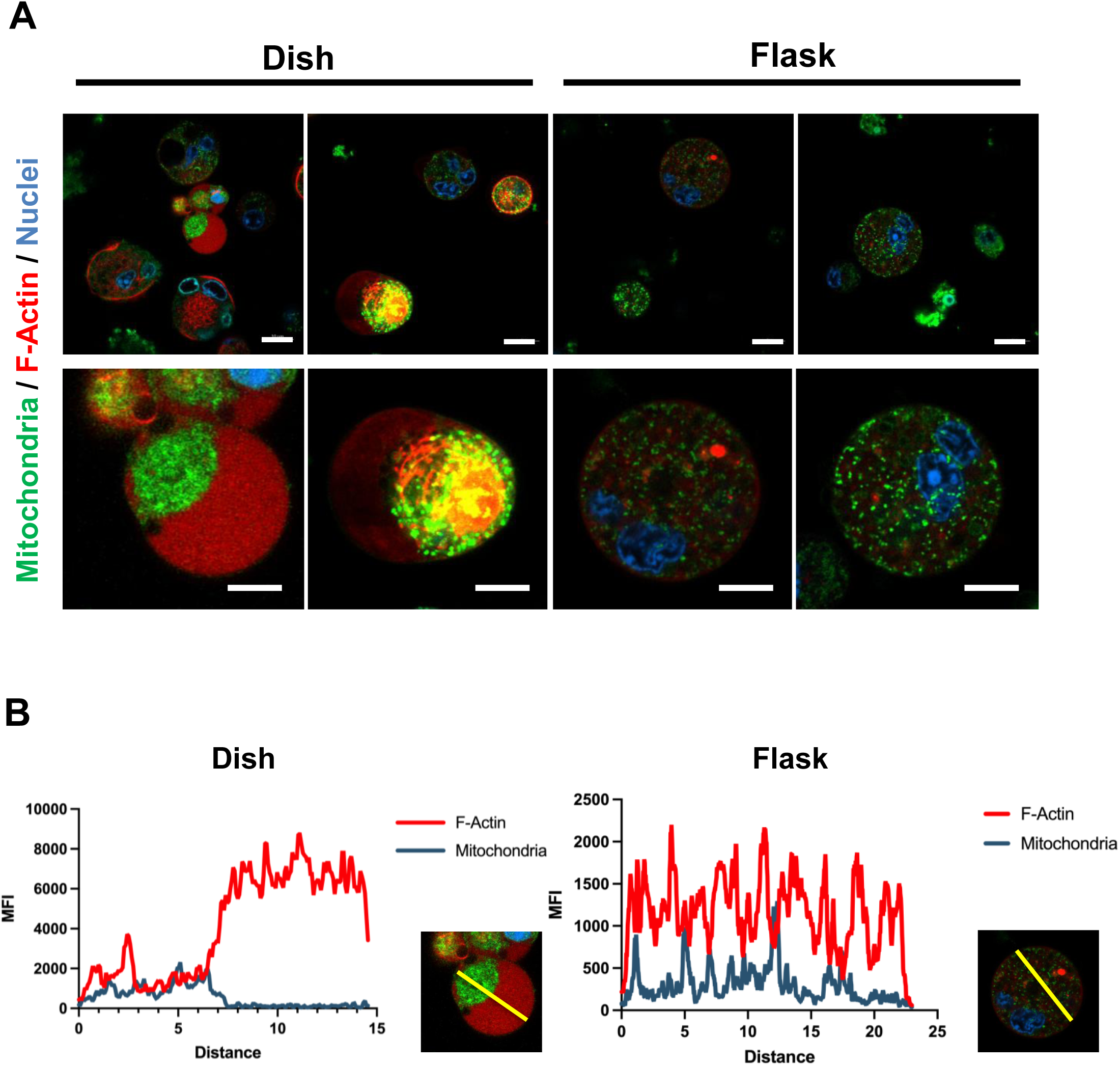
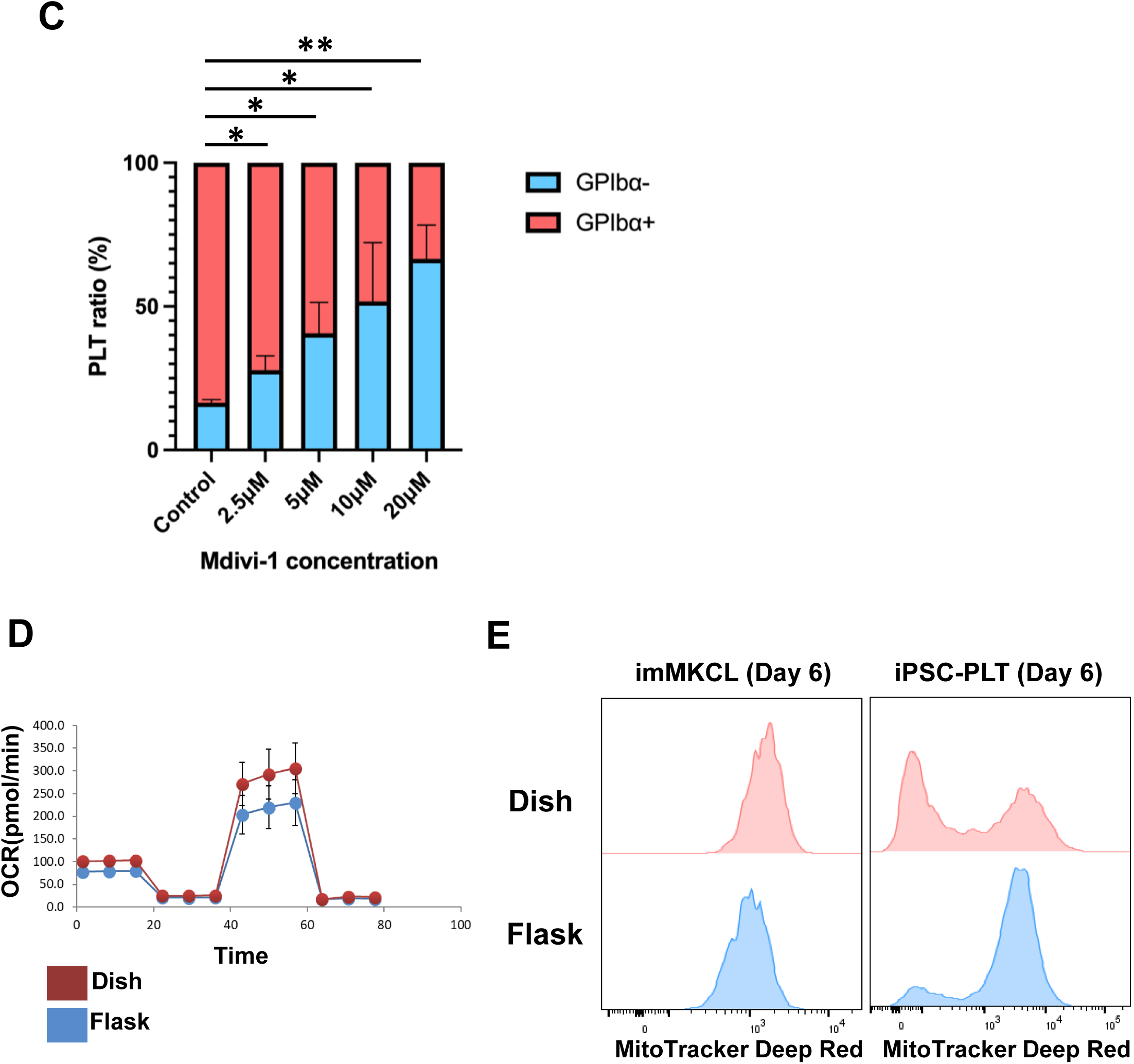
**Turbulent culture conditions promote uniform mitochondrial distribution in imMKCLs** (A, B) Confocal imaging and fluorescence intensity analysis of day 4 maturing imMKCLs cultured under static (dish) or turbulent (flask) conditions. (A) Representative confocal images of day 4 maturing imMKCLs. Top panels show original images (scale bars, 10 µm); bottom panels show magnified views (scale bars, 5 µm) of cells cultured in a dish (left) or flask (right). Mitochondria were stained with MitoTracker Green (green), F-actin with SPY555-FastAct (red), and nuclei with Hoechst 33342 (blue). (B) Fluorescence intensity profiles of F-actin (red) and mitochondria (blue) along the central axis (yellow line in A, magnified images). (C) Quantification of the percentage of GPIbα⁺ (red) and GPIbα⁻ (blue) platelets on day 6 following Mdivi-1 treatment. Data are shown as mean ± SEM (n = 3). Statistical significance was determined using an unpaired two-tailed t-test (*p < 0.05; **p < 0.01). (D) Oxygen consumption rate (OCR) of day 4 maturing imMKCLs cultured in a dish (red) or flask (blue), measured using the Seahorse Flux Analyzer. (E) Representative histograms showing mitochondrial membrane potential (MitoTracker Deep Red) in day 6 imMKCLs (left) and iPSC-platelets (right) cultured in a dish (top, pink) or flask (bottom, blue).

### Turbulence regulates actin dynamics to produce high-quality platelets

The actin cytoskeleton is known to play a crucial role in the distribution of mitochondria during cell division (Moore et al., 2021). To determine whether actin contributes to the uniform mitochondrial distribution in imMKCLs, we examined cytoskeletal dynamics under different culture conditions. Interestingly, there were no significant differences in the expression of cytoskeleton-related genes (Fig. 5, A–C) or total actin protein levels (Fig. 5, D and E) between dish and flask cultures. However, from day 4 onwards, we observed a marked, turbulence-dependent reduction in F-actin-specific signals in turbulent cultures compared to static dish cultures (Fig. 5 F).

**Figure 5.**
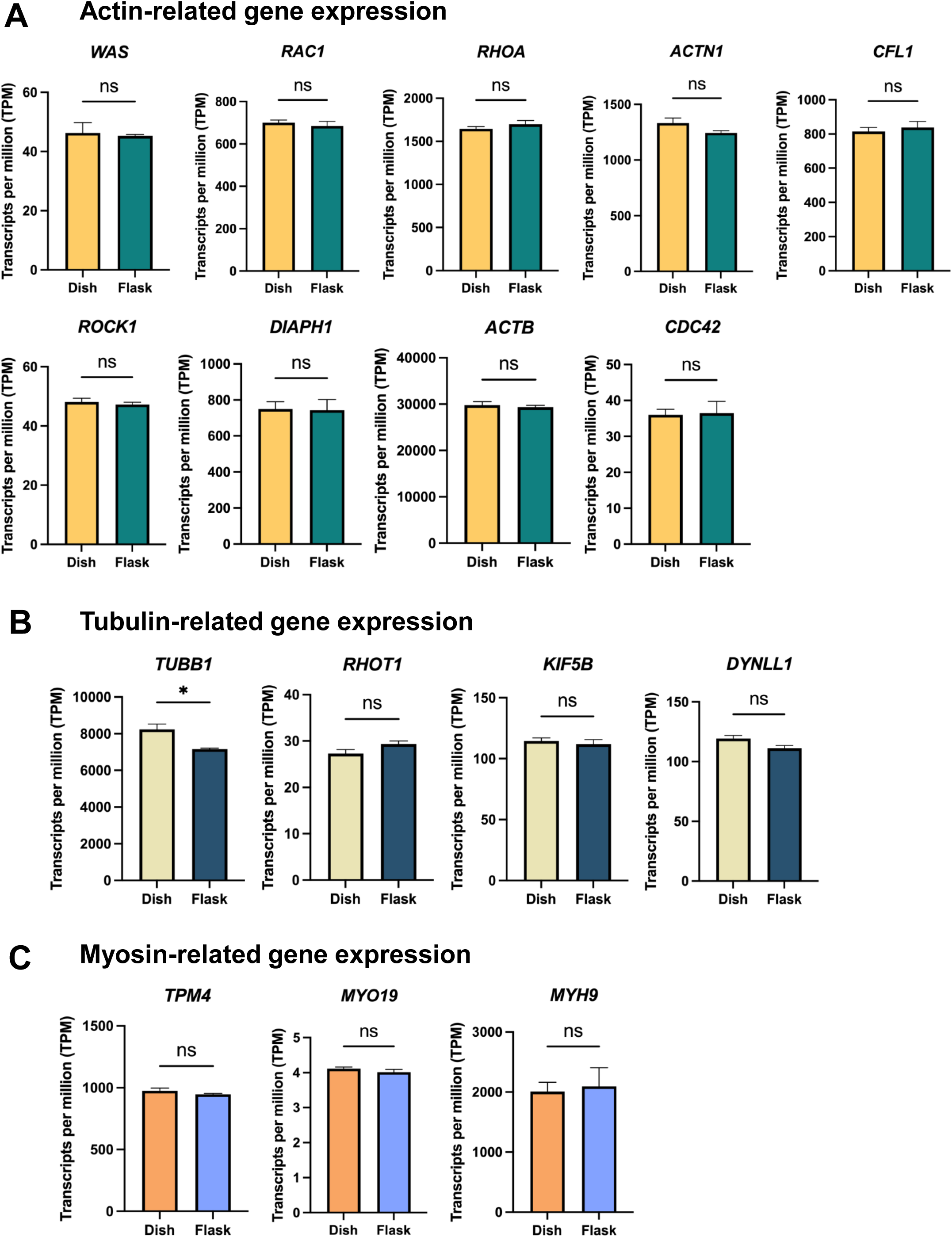
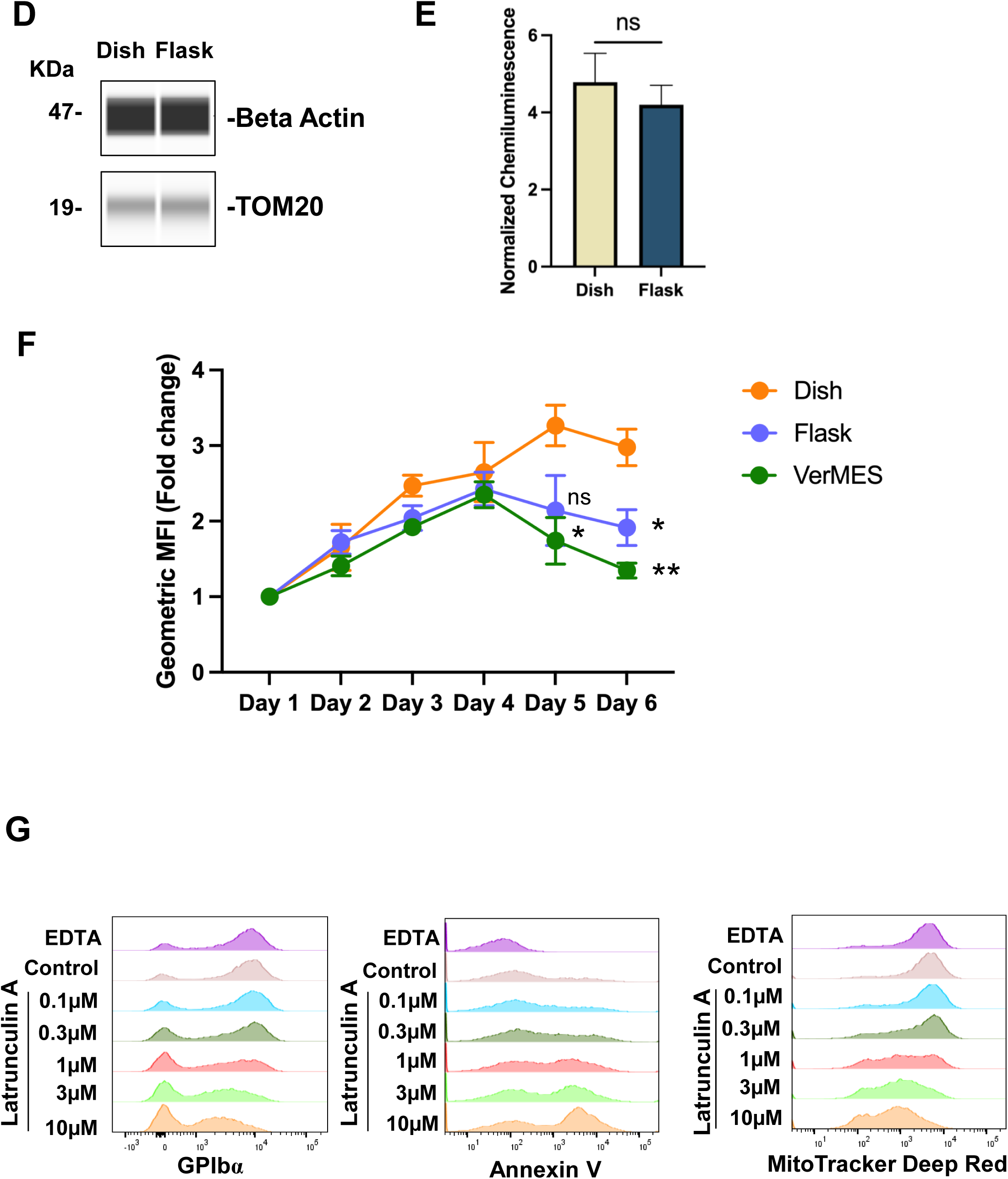
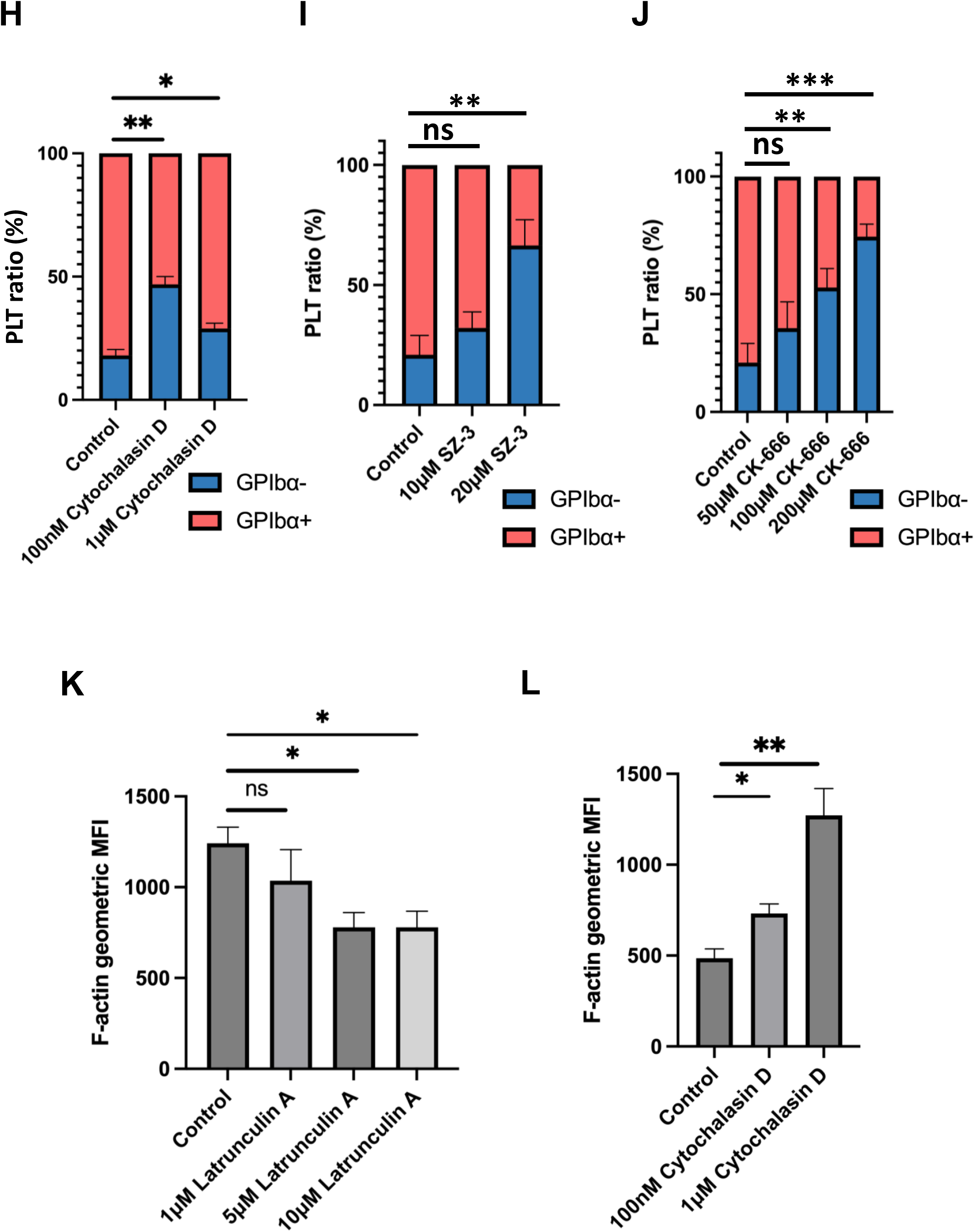
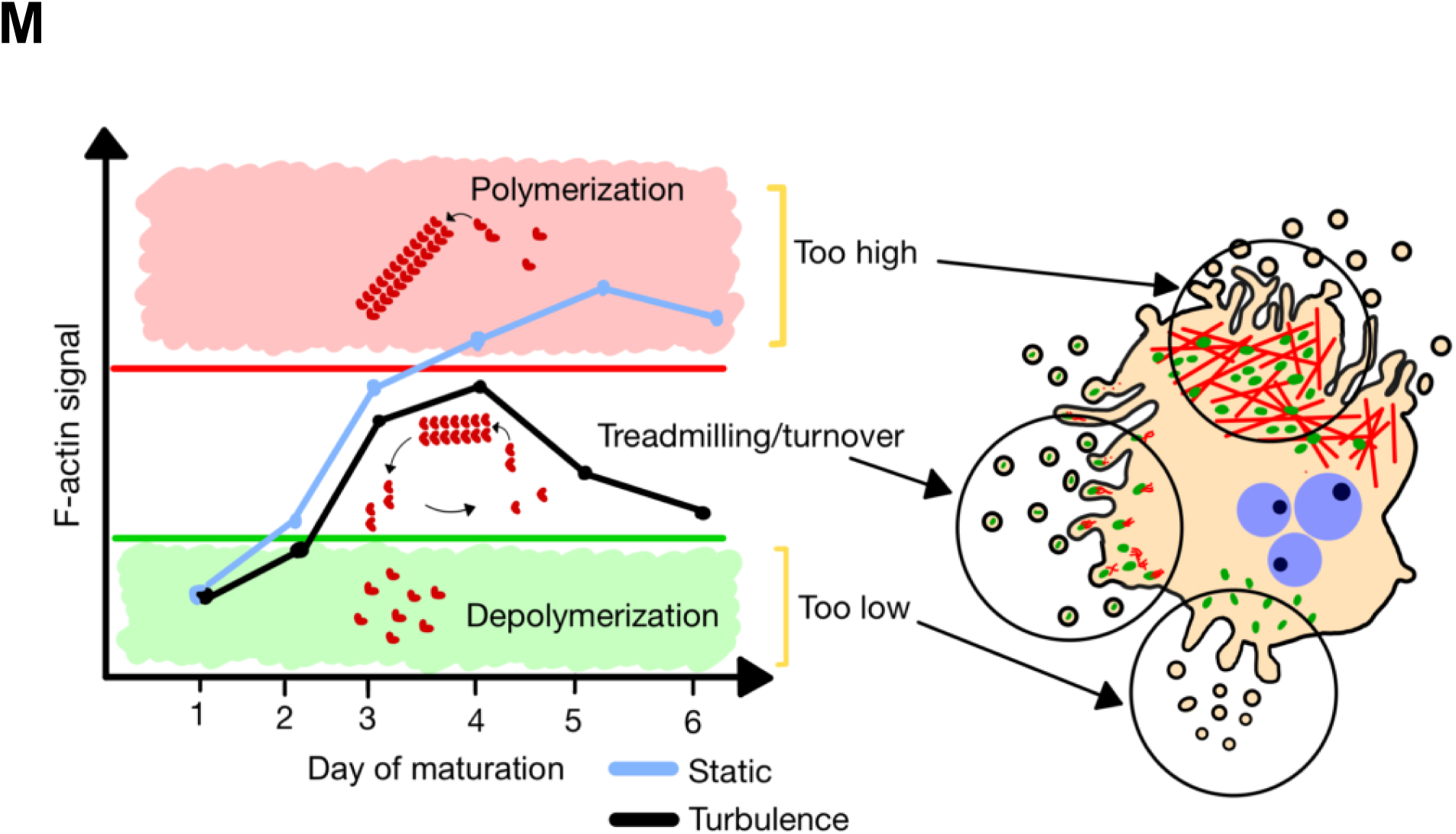
Turbulence regulates actin dynamics to promote high-quality platelet production. (A–C) Transcript levels of cytoskeletal genes related to actin (A), tubulin (B), and myosin (C) in day 4 maturing imMKCLs cultured in a dish or flask. Gene expression is shown as transcripts per million (TPM). Data are presented as mean ± SEM (n = 3). Statistical analysis was performed using an unpaired two-tailed t-test (ns, not significant). (D, E) Protein levels of β-actin normalized to TOM20 in day 5 maturing imMKCLs cultured in a dish (orange) or flask (purple). (D) Representative Simple Western (Wes) images. (E) Quantification of β-actin expression. Data are presented as mean ± SEM (n = 3). Statistical analysis was performed using an unpaired two-tailed t-test (ns, not significant). (F) Fold change in F-actin levels (SPY555-FastAct signal) from day 1 to day 6 in maturing imMKCLs cultured in a dish (orange), flask (purple), or VerMES (green). Data are presented as mean ± SEM (n = 3). Statistical comparisons were performed independently on day 5 and day 6 between dish and flask and between dish and VerMES using unpaired two-tailed t-tests (ns, not significant; *p < 0.05; **p < 0.01). (G) Representative histograms showing surface GPIbα expression (left), phosphatidylserine exposure (annexin V binding, middle), and mitochondrial membrane potential (MitoTracker Deep Red, right) in day 6 iPSC-derived platelets (iPSC-PLTs) cultured with Latrunculin A at 0.1 µM (dark green), 0.3 µM (lime green), 1 µM (orange), 3 µM (blue), or 10 µM (red). EDTA-treated (purple) and untreated control (mauve) samples are shown at the top. (H–J) Quantification of the percentage of GPIbα⁺ (red) and GPIbα⁻ (blue) platelets on day 6 following treatment with: (H) 100 nM or 1 µM Cytochalasin D, (I) 10 µM or 20 µM SZ-3, or (J) 50 µM, 100 µM, or 200 µM CK-666. Data are presented as mean ± SEM (n = 3). Statistical analysis was performed using an unpaired two-tailed t-test (ns, not significant; **p < 0.01; ***p < 0.001; ****p < 0.0001). (K, L) Quantification of F-actin levels in day 5 imMKCLs using SPY555-FastAct following actin-targeting compound treatment. Geometric mean fluorescence intensity is shown for cells treated on day 3 with (K) 1, 5, and 10 µM Latrunculin A, or (L) 100 nM and 1 µM Cytochalasin D. (M) Schematic model illustrating the relationship between F-actin levels, the polymerization/depolymerization state of intracellular actin, and their effects on platelet production in maturing imMKCLs.

**Figure 6.**
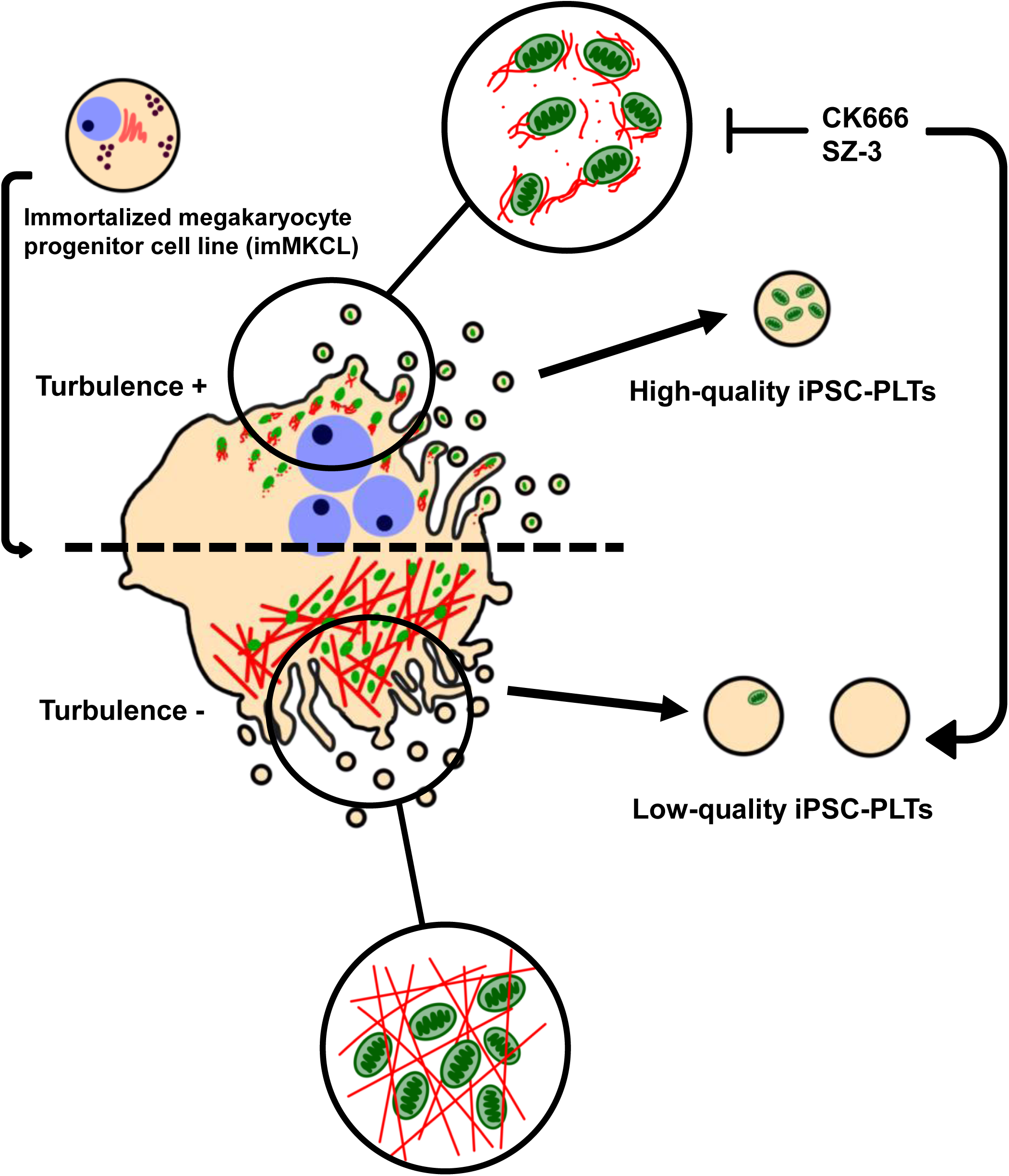
**Graphical summary** Turbulence reduces actin (red) polymerization during imMKCL maturation, facilitating efficient incorporation of mitochondria (green) into the resulting iPSC-platelets. This process is disrupted by actin polymerization inhibitors (CK-666, SZ-3), leading to the production of low-quality platelets similar to those generated under static culture conditions.

To explore how reduced F-actin levels—detected by SPY555-FastAct staining—might relate to mitochondrial delivery, we first investigated the effects of Latrunculin A, a monomeric actin-binding molecule that prevents actin polymerization. Treatment with Latrunculin A resulted in a dose-dependent decrease in both GPIbα expression and mitochondrial membrane potential (MMP), accompanied by increased annexin V binding as well as decreased F-actin levels in imMKCLs (Fig. 5, G and K). Conversely, treatment with Cytochalasin D, which protects actin filament barbed ends, but not pointed ends, from actin polymerization, significantly enhanced F-actin levels in imMKCLs but still led to diminished GPIbα expression in iPSC-platelets (Fig. 5, H and L). These results suggest that an intermediate, dynamically relaxed actin structure— characterized by moderate F-actin levels (Fig. 5 F)—may be optimal for functional platelet production.

To further assess whether actin polymerization-depolymerization dynamics influence GPIbα expression in iPSC-platelets, we evaluated the roles of cofilin, a key mediator of actin depolymerization, and the Arp2/3 complex, a regulator of actin branching (reference). Inhibition of cofilin using SZ-3, or the Arp2/3 complex using CK-666, also resulted in reduced GPIbα expression (Fig. 5, I and J), further supporting the importance of actin dynamics in platelet production.

Collectively, these findings highlight the pivotal role of turbulence in modulating actin dynamics and mitochondrial distribution, both essential for generating functional iPSC-platelets. We therefore propose that the dynamic turnover of actin filaments—through a process of balanced polymerization and depolymerization, known as treadmilling—represents a critical mechanism for enabling optimal mitochondrial delivery and efficient iPSC-platelet production (Fig. 5 K).

## DISCUSSION

In this study, we sought to elucidate the underlying mechanisms by which turbulent flow facilitates the generation of GPIbα⁺ iPSC-platelets from imMKCLs, as previously described (Ito et al., 2018). To this end, we conducted comparative analyses under turbulent flow and static culture conditions (Fig. 1 A; Fig. S1 A). These experiments revealed that both the absolute platelet number and proportion of GPIbα⁺ iPSC-platelets were significantly increased under turbulent conditions (Fig. 1, B–D). Further characterization confirmed the high quality of these platelets, as demonstrated by assays assessing GPIIb/IIIa activation and P-selectin (CD62P) exposure (Fig. S1 B), annexin V binding (Fig. S1 C), Calcein-AM retention (Fig. 1 F) (Moreau et al., 2016; Zhao et al., 2023), and uptake of fluorescently labeled Factor V (Fig. S1, J–L) (Sim et al., 2017). GPIbα expression levels were correlated with other activation-related markers, such as PAR1 and GPVI (Fig. S1 D). These markers are also known to undergo ADAM-mediated shedding, similar to GPIbα, suggesting that GPIbα⁻ platelets may be more prone to shedding under whole-cell conditions (Ludeman et al., 2004; Gardiner et al., 2007).

Notably, GPIbα⁺ iPSC-platelets retained functional mitochondria, as evidenced by preserved mitochondrial signals, maintained membrane potential as well as robust ROS and ATP production (Fig. 1 E, Fig. 2, A and C). Using peripheral blood-derived human platelets, we confirmed that prolonged storage at 22°C or 37°C led to a decline in mitochondrial membrane potential, accompanied by reduced GPIbα expression and increased phosphatidylserine (PS) exposure (Fig. S2, D and E).

These findings suggest that turbulent flow enhances the efficient incorporation of mitochondria into platelet particles. A culture-switch experiment further demonstrated that introducing turbulence from day 4 onward—corresponding to the late maturation stage of imMKCLs—is critical for increasing GPIbα⁺ iPSC-platelet yields (Fig. 1, G and H), indicating that mitochondrial allocation into platelets is regulated during late-stage megakaryocyte maturation and subsequent platelet release (Fig. S1 F). Concomitantly, we observed a turbulence-specific reduction in F-actin levels from day 4, prompting the hypothesis that turbulence modulates F-actin dynamics to facilitate mitochondrial distribution, a point we explored further below.

Previous reports have shown that GPIbα is subject to ectodomain shedding via ADAM17, reducing its surface expression on both iPSC-and primary platelets (Bergmeier et al., 2004; Nishikii et al., 2008). However, the upstream regulatory mechanisms have remained elusive. In this study, we demonstrate that GPIbα⁺ iPSC-platelets maintain ATP-dependent flippase activity, which actively suppresses PS externalization. In contrast, impaired mitochondrial function leads to PS exposure and subsequent ADAM17-mediated GPIbα shedding.

Importantly, scramblase activity was not elevated in GPIbα⁻ iPSC-platelets, suggesting that PS exposure in this population arises solely from diminished flippase activity—a departure from earlier findings implicating Ca²⁺-dependent scramblase TMEM16F in PS externalization (Fujii et al., 2015). Ionomycin stimulation failed to elicit Ca²⁺ influx in GPIbα⁻ iPSC-platelets (Fig. 3 D), indicating a deficiency in the calcium regulation, in contrast to intact peripheral platelets. Supporting this, peripheral blood platelets subjected to long-term storage at 22°C exhibited increased scramblase and decreased flippase activity (Fig. S3 B).

While our previous work demonstrated that the ADAM17 inhibitor KP457 enhances the production of GPIbα⁺ iPSC-platelets (Hirata et al., 2017), our current findings suggest that mitochondrial dysfunction is an upstream event regulating GPIbα shedding. Importantly, GPIbα expression itself did not affect mitochondrial function, as shown in GPIX knockout (KO) imMKCLs (Fig. 2, E and F). Furthermore, KP457 did not influence the number or function of resulting platelets (Fig. S1, G–I). Despite prior use of KP457 in culture, our data put into question its necessity for future iPSC-platelet manufacturing protocols.

In cells, including platelets, intracellular ATP production is supported not only by mitochondrial oxidative phosphorylation but also by glycolysis. While earlier studies reported that glycolysis is the predominant ATP source in platelets (Akkerman and Holmsen, 1981), recent advances in analytical techniques have shown that mitochondrial ATP production may actually be dominant, with oxidative phosphorylation accounting for approximately 75% of total ATP generation (Jedlička et al., 2021). These findings suggest that mitochondrial dysfunction can cause intracellular ATP depletion to an extent sufficient to impair flippase activity and ultimately trigger GPIbα shedding, as demonstrated by treatment with the depolarizing agent ionomycin (Fig. 3 E).

Mechanistically, turbulence was found to optimize F-actin dynamics by finetuning the balance between polymerization and depolymerization, thereby ensuring uniform mitochondrial distribution within maturing megakaryocytes and promoting efficient platelet biogenesis. The reduction in F-actin levels under turbulent conditions during the late maturation stage (as noted earlier) highlights this critical window for enhanced GPIbα⁺ iPSC-platelet generation. To validate this, we treated cultures with compounds targeting F-actin (Latrunculin A, Cytochalasin D), cofilin (SZ-3), the Arp2/3 complex (CK-666), and Drp1 (Mdivi-1). Each of these interventions led to a dose-dependent decline in GPIbα⁺ iPSC-platelet output, although mitochondrial membrane potential (MMP) levels remained relatively unchanged.

Interestingly, previous work has implicated actin polymerization in promoting GPIbα shedding (Zhou et al., 2022). While Cytochalasin D treatment paradoxically increased F-actin levels, Latrunculin A reduced both F-actin content and GPIbα expression. Actin filaments were visualized using a fluorescent derivative of the cell-permeable F-actin-stabilizing compound, jasplakinolide (Milroy et al., 2012; Merino et al., 2020). As jasplakinolide stabilizes actin filaments, increases in the F-actin signal should be interpreted with caution (Pospich et al., 2020; Merino et al., 2020). Nevertheless, the data support a critical role for actin filament dynamics in mitochondrial distribution and GPIbα expression. These findings suggest that GPIbα shedding is not simply dictated by polymerized actin quantity but rather by the dynamic equilibrium of actin turnover. Consistent with this, image analyses revealed uniform mitochondrial distribution in imMKCLs exposed to turbulent flow. Given that actin treadmilling is crucial for organelle distribution from mother to daughter cells (Moore et al., 2021) and neuronal mitochondrial transport relies predominantly on microtubules, our results instead emphasize an actin-centric mechanism in megakaryocytes. Together, these data support a model in which actin treadmilling—characterized by continuous polymerization and depolymerization—governs the even allocation of mitochondria and is fundamental to the production of functional platelets.

While this study underscores the pivotal role of turbulence in regulating F-actin dynamics and mitochondrial distribution, the precise mechanotransductive link between turbulent forces and cytoskeletal modulation in megakaryocytes remains to be further defined. Future work utilizing confocal and super-resolution microscopy, in conjunction with analysis of specific actin-binding proteins, will be critical for dissecting actin dynamics in ex vivo megakaryocyte cultures. Further clarification of the roles of cofilin and the Arp2/3 complex in this context is also warranted. Finally, identifying the extracellular cues that influence intracellular actin organization will provide valuable insights for refining iPSC-platelet manufacturing strategies.

In summary, our findings reveal that turbulent flow is a key regulator of actin-dependent mitochondrial segregation, a process central to generating high-quality iPSC-platelets. These insights not only advance our understanding of platelet biogenesis but also offer practical avenues to improve the scalability and quality of iPSC-platelet manufacturing.

## Materials and Methods

### Cells

imMKCL Clone 17, originally established by the Eto group at Kyoto University, was used in all experiments except those related to *GP9* knockout (GPIX-KO). GPIX-KO cells were generated by the Freson group at KU Leuven from clone 7 imMKCL, using the CRISPR-Cas9 system to introduce a frameshift variant in the *GP9* gene, creating a homozygous deletion. Immunoblot analysis confirmed the absence of GPIX expression and GPIbα in GPIX-KO cells (De Wispelaere et al., in revision, *HemaSphere*).

### Culture medium

The basal medium consisted of IMDM (Sigma-Aldrich, #I3390) supplemented with 2 mM L-glutamine (Thermo Fisher Scientific, #25030-081), insulin-transferrin-selenium (Thermo Fisher Scientific, #41400-045), 50 µg/mL ascorbic acid (Sigma-Aldrich, #A4544), and 450 µM 1-thioglycerol (Sigma-Aldrich, #M6145). For DOX-ON cultures, the basal medium was further supplemented with 15% fetal bovine serum (FBS; Sigma-Aldrich, #172012), 50 ng/mL animal component-free recombinant human stem cell factor (SCF; Wako, #193-15513), 200 ng/mL TA-316 (Nissan Chemical), and 5 µg/mL doxycycline (Clontech, #631311). For DOX-OFF cultures, the basal medium was supplemented with human plasma (Japan Blood Products Organization), 1 IU/mL enoxaparin sodium (Kaken Pharmaceutical Co., Ltd.), 50 IU/mL Cleactor Intravenous Infusion (Eisai Co., Ltd., #3959407D2034), 50 ng/mL SCF, 200 ng/mL TA-316, 0.75 µM SR-1 (Calbiochem) or 0.5 µM GNF-351 (Calbiochem), and 10 µM Y-27632 (Wako) or 0.5 µM Y-39983 (MedChemExpress).

### Cell culture

#### DOX-ON proliferation culture

The expansion culture (DOX-ON stage) was performed as previously described (Ito et al., 2018). Briefly, cells were passaged twice a week, maintaining cell densities between 1 × 10⁵ – 2 × 10⁶ cells/mL throughout the culture period. For all experiments except those using the VerMES bioreactor, DOX-ON cultures were conducted in 10-cm culture dishes (Thermo Scientific, #150466). For larger-scale experiments using the VerMES bioreactor, cell expansion was achieved through sequential upsizing as follows: starting with 10-cm static dishes, followed by 125 mL and 500 mL Corning Erlenmeyer cell culture flasks (Sigma-Aldrich, #431143 and #431145) under shaking conditions using a Lab-Therm shaker (Kuhner), and finally scaled up to a 10 L CELLBAG™ operated with the ReadyToProcess WAVE 25 system (Cytiva, #CB0010L10-03) under rocking motion.

#### DOX-OFF differentiation culture

For platelet production following imMKCL maturation (Dox-OFF stage), approximately 1 – 3 × 10⁵ cells/mL were cultured for 6 days under one of the following conditions: in 10-cm culture dishes (Thermo Scientific, #150466), in 125 mL Corning Erlenmeyer cell culture flasks (Sigma-Aldrich, #431143) under shaking conditions using a Lab-Therm shaker (Kuhner), or in a VerMES bioreactor (SATAKE Co., Ltd., Saitama, Japan). The VerMES bioreactor was equipped with vertically oscillating two-stage mixing blades, and the optimal operating conditions for the 8 L VerMES were a stroke of 40 mm and a speed of 150 mm/s.

#### Human platelets

Donor-derived human platelets were provided by the Japanese Red Cross Society. Human platelets and plasma were used in compliance with the Guidelines on the use of Donated blood in R&D, etc., from the Ministry of Health, Labour and Welfare of Japan.

#### Flow cytometric analysis

Flow cytometric analysis was performed using a FACSLyric cytometer (BD Biosciences) for the experiments described below (platelet count, annexin V binding assay, platelet activation, flippase and scramblase activity, calcium assay, factor V assay, Calcein-AM assay, mitochondrial measurements, and F-actin detection). The following antibodies and reagents were used: anti-human CD41-APC (BioLegend, #303710), anti-human CD41-PE (BioLegend, #303706), anti-human CD42b-PE (BioLegend, #303906), anti-human CD42b-PerCP (BioLegend, #303910), anti-human CD42a-FITC (eBioscience, #11-0428-42), anti-human GPVI-Brilliant Violet 421 (BD Biosciences, #743941), anti-human CD62P-Brilliant Violet 421 (BioLegend, #304926), PAC-1-FITC (BD Biosciences, #340507), and Annexin V-FITC (BD Biosciences, #556419). All flow cytometry data were analyzed using FlowJo v10 (BD Biosciences).

#### Platelet count

The number of platelet particles per imMKCL cell was defined as the total count of CD41⁺ iPSC-platelets or CD41⁺CD42b⁺ iPSC-platelets on the indicated day, divided by the number of imMKCL cells on day 0 of the Dox-OFF culture. Platelet counts were determined using Trucount Tubes (BD Biosciences, #340334).

#### Annexin V binding assay

Staining with FITC-conjugated annexin V, together with anti-CD41 and anti-CD42b antibodies, was performed in the annexin V binding buffer (BD Biosciences, #556454) supplemented with 1 μM CaCl₂. For positive and negative controls, 20 µM ionomycin (Wako, #091-05833) or EDTA (Nacalai Tesque, #06894-14) were added to each sample, respectively, prior to staining.

#### Platelets activation

Platelet activation was evaluated by measuring PAC-1 binding and P-selectin (CD62P) expression, following the protocol described in our previous study (Sone et al., 2021). Briefly, culture suspensions of iPSC-platelets were stimulated with or without 0.2 µM phorbol 12-myristate 13-acetate (PMA) or a combination of 100 µM adenosine diphosphate (ADP) and 40 µM thrombin receptor-activating peptide 6 (TRAP-6). Cells were then incubated with anti-CD62P, anti-human CD41a, and PAC-1 antibodies for 30 minutes at room temperature. After incubation, samples were diluted in HEPES-Tyrode buffer (10 mM HEPES, 15 mM NaHCO₃, 138 mM NaCl, 5.5 mM glucose, 2.6 mM KCl, 1 mM MgCl₂, and 2 mM CaCl₂) and analyzed by flow cytometry.

#### Flippase and scramblase activity

Flippase and scramblase activity was measured as previously described (Miyata and Segawa, 2022). Briefly, iPSC-platelets were stained with 0.5 µM 18:1-06:0 NBD-PS (Merck, #810194P) for 20 minutes or 0.5 µM 16:0-06:0 NBD-PC (Merck, #810130P) for 60 minutes, together with anti-CD41 and anti-CD42b antibodies. After staining, 2% BSA (prepared by diluting a 10% stock solution; Merck, #A1595) was added to remove non-specific binding. To confirm staining specificity, 60 µM ionomycin was added during NBD-PC or NBD-PS staining.

#### Calcium assay

iPSC-platelets were stained with 3 µL Fluo-4 AM (Dojindo Molecular Technologies, Inc., #F312) together with anti-CD41 and anti-CD42b antibodies for 20 minutes. To assess calcium influx in response to stimulation, 2 µM ionomycin was added immediately prior to flow cytometric analysis.

#### Factor V assay

Human factor V (Haematologic Technologies Inc., #HCV-0100) was fluorescently labeled using the Fluorescent Protein Labeling Kit (Thermo Fisher Scientific, #A10235) prior to its addition to imMKCL cultures on day 3 of the Dox-OFF stage. The incorporation of factor V into the produced platelets was evaluated on day 6.

#### Calcein-AM assay

Calcein-AM (FUJIFILM Wako Pure Chemical Corporation, #341-07901) uptake by iPSC-platelets was assessed on day 6 of Dox-OFF cultures. Platelets were stained with 2 µM Calcein-AM together with anti-CD41 and anti-CD42b antibodies for 30 minutes.

#### Mitochondrial measurements

The reagents and their concentrations used in these experiments were as follows: 100 µM BioTracker ATP-Red Live Cell Dye (Merck, #SCT045) for ATP content; 1 µM CellROX Deep Red Flow Cytometry Assay Kit (Thermo Fisher Scientific, #C10491) for reactive oxygen species (ROS) production; 100 nM MitoTracker Deep Red (Thermo Fisher Scientific, #M22426) or the MT-1 MitoMP Detection Kit (Dojindo Molecular Technologies, Inc., #MT13) for mitochondrial membrane potential; 1:10,000 dilution of PicoGreen (Thermo Fisher Scientific, #P11495) for mitochondrial DNA detection; and 50 nM MitoTracker Green (Dojindo Molecular Technologies, Inc., #MT10) for mitochondrial presence. Platelets or imMKCLs were stained with each reagent and antibodies against CD41 and CD42b for 30 minutes.

#### F-actin detection

For F-actin-specific detection, platelets or imMKCLs were stained with SPY555-FastAct (Cytoskeleton, Inc., #CY-SC205) together with an anti-CD41 antibody for 2 hours.

#### Metabolic flux analysis

imMKCL cells were seeded into wells of an XF96 microplate pre-coated with 0.1% poly-L-lysine to measure the oxygen consumption rate (OCR) using an Agilent Seahorse XF Pro analyzer. The assay was performed with the Seahorse XF Cell Mito Stress Test Kit (Agilent Technologies, #103015-100) and XF RPMI Medium, pH 7.4 (Agilent Technologies, #103576-100). After seeding, the plate was centrifuged at 200 × *g* for 1 minute to form a monolayer, then equilibrated for 40 minutes at 37 °C in a CO₂ incubator before starting the assay. The Cell Mito Stress Test protocol included sequential injections of oligomycin (1 µM), FCCP (2 µM), and a mixture of rotenone and antimycin A (1 µM each). For each injection, four measurement cycles were performed, consisting of 3 minutes of mixing followed by 3 minutes of measurement. OCR data were analyzed using Wave 2.6 software (Agilent Technologies).

#### Protein Quantification Using Wes System^TM^

Protein levels were analyzed using the Wes^TM^ automated capillary electrophoresis-based immunoassay system (ProteinSimple). Cells were lysed in RIPA buffer supplemented with protease and phosphatase inhibitors, and protein concentrations were measured using the BCA Protein Assay Kit (Thermo Fisher Scientific). The following antibodies were used: anti-β-actin (Cell Signaling Technology, #4970), anti-TOM20 (Abcam, #ab186735), and Anti-Rabbit Detection Module Chemiluminescence (ProteinSimple, #DM-001). Equal amounts of total protein, blocking reagent, primary and secondary antibodies, and chemiluminescent substrate were loaded into the Wes^TM^ assay plate according to the manufacturer’s instructions.

Chemiluminescent signals were acquired and quantified using Compass for Simple Western software (ProteinSimple). β-actin levels were normalized to TOM20.

#### Electron Microscopy

Platelet pellets were fixed with a mixture of 0.5% glutaraldehyde and 2% paraformaldehyde in 0.1 M phosphate buffer (pH 7.4) for 60 minutes at 4 °C. After washing with phosphate buffer, samples were post-fixed with 1% osmium tetroxide in phosphate buffer for 60 minutes on ice. Following dehydration, pellets were infiltrated with and embedded in epoxy resin. Ultrathin sections (60–80 nm thick) were cut, stained with 2% uranyl acetate in 70% methanol and Reynolds’ lead citrate, and examined using a transmission electron microscope operating at 80 kV (HT-7700; Hitachi, Tokyo, Japan).

#### Live-cell imaging

For live-cell imaging using confocal microscopy, u-Dish 35 mm Quad ibiTreat sterilized dishes (Nippon Genetics, #ib80416), pre-coated with 0.1% poly-L-lysine, were used. Imaging was performed using a ZEISS LSM 900 confocal microscope equipped with Airyscan 2 and a 63×/1.40 numerical aperture oil-immersion objective. The following reagents were used for staining imMKCLs or platelets: APC-conjugated anti-human CD41 antibody, PerCP-conjugated anti-human CD41 antibody, MitoTracker Green, SPY555-FastAct, Plasmem Bright Green (Dojindo Laboratories, #P504), and Hoechst 33342 (Thermo Fisher Scientific, #62249). Confocal images were acquired using the same system.

#### Small molecule compound treatments

Mdivi-1 (Sigma-Aldrich, #M0199-5MG) was added on day 0 of the DOX-OFF culture to inhibit mitochondrial fission. Latrunculin A (FUJIFILM Wako Pure Chemical Corporation, #125-04363), Cytochalasin D (Cayman Chemical, #11330), SZ-3 (MedKoo Biosciences, #208333), and CK-666 (Selleck, S7690) were added on day 3 of the Dox-OFF culture. Various ADAM inhibitors were used as follows: 15 µM KP-457 (Kaken Pharmaceutical), 10 µM GI254023X (Selleck, #S8660), 100 µM TAPI-1 (Selleck, #S7434), and 10 µM GM6001 (Ilomastat; Selleck, #S7157). For all experiments, dimethyl sulfoxide (DMSO) was added to control wells and used as the solvent for compound dilution.

#### Mitochondrial mAG overexpression

The CoralHue^TM^ mitochondria-targeted monomeric Azami-Green (pMT-mAG1) plasmid was provided by RIKEN BRC through the National BioResource Project of MEXT, Japan (Cat. #RDB18311). A *piggyBac* transposon vector system was used to establish stable mitochondrial labeling in imMKCLs. The gene of interest (GOI), encoding CoralHue mitochondria-targeted monomeric Azami-Green, was amplified from pMT-mAG1 (Karasawa et al., 2003) via PCR. The amplified GOI was inserted into the backbone vector using the In-Fusion HD Cloning Kit (Clontech, #639649).

#### Bulk RNA-sequencing analysis

Total RNA was extracted from cultured imMKCLs on DOX-OFF day 4 using the RNeasy Plus Micro Kit (Qiagen, #74034). For each sample, up to 10 ng of total RNA was processed using the SMART-Seq v4 Ultra Low Input RNA Kit for Sequencing (Clontech, #634890).

Complementary DNA (cDNA) was fragmented using an S220 Focused-ultrasonicator (Covaris), and the cDNA library was prepared and amplified with the NEBNext Ultra DNA Library Prep Kit for Illumina (New England Biolabs, #E7370L). Library size distribution was evaluated using a Bioanalyzer with the Agilent High Sensitivity DNA Kit. Sequencing was performed on an Illumina NextSeq 500. FASTQ files were generated using *bcl2fastq* v2.20. Adapter sequences and low-quality bases were trimmed from raw sequencing reads using *cutadapt* v4.1 (Kechin et al., 2017). Trimmed reads were aligned to the human reference genome (hg38) using *STAR* v2.7.10a (Dobin et al., 2013) with the GENCODE annotation file (release 32, GRCh38.p13) (Frankish et al., 2019). Uniquely mapped reads were quantified at the gene level using *htseq-count* v2.0.2 (Anders et al., 2015) with the same GENCODE GTF file. Gene expression levels were calculated and expressed as transcripts per million (TPM).

#### Statistical analysis

Statistical analysis was performed using GraphPad Prism 10 (GraphPad Software, La Jolla, CA). Data are expressed as the mean ± standard error of the mean (SEM). To determine statistical significance, an unpaired two-tailed Student’s t-test was used. Details of sample sizes, statistics, and statistical significance are indicated in each figure legend.

## Data and code availability

RNA-seq data have been deposited in the NCBI Gene Expression Omnibus (GEO) under accession number **GSE298561**.

## Acknowledgements

We thank Dr. Kelvin Hui for the critical reading of this manuscript. We also appreciate Keiko Yamamoto for supporting the immunocytochemistry analysis and cell culture and the Japanese Red Cross for providing donor-derived platelets.

This work was supported by the Japan Agency for Medical Research and Development (AMED) under grant numbers JP20bm0704051 (K.E.) and JP23bm1323001 (S.N. and K.E.); Otsuka Pharmaceutical Co. Ltd. (to K.E.); the CiRA Foundation Fund (to K.E.); the Canon Foundation (to K.E.); Grants-in-Aid for Scientific Research from the Japan Society for the Promotion of Science (JSPS) (KIBAN S, 21H05047 to K.E.; and Young Scientists, 22K18169 to S.N.); the JST FOREST Program (JPMJFR225K to S.N.); JST CREST (JPMJCR24B3 to S.S. and K.E.); the New Energy and Industrial Technology Development Organization (NEDO, JPNP23028 to K.E.); and by KU Leuven grants including the BOF grant (C14/23/121 to K. Freson) and the FWO grant (G072921N to K. Freson).

## Author contributions

E.N. and Y.H. conducted most experiments and analyzed data. K. Freson., K.D.W., and C.T. generated GPIX-KO clone 7 imMKCLs. A.S. conducted the platelet transmission electron microscopy and analyzed data. K. Fujio. constructed the vector system and analyzed related results. S.N. and S.S. supervised the experimental design. E.N., Y.H., and K.E. wrote the manuscript. K.E. edited the manuscript.

## References

1. Ajanel, A., R.A. Campbell, and F. Denorme. 2023. Platelet mitochondria: the mighty few. Curr. Opin. Hematol. 30:167–174. doi:10.1097/MOH.0000000000000772.

2. Akkerman, J.W., and H. Holmsen. 1981. Interrelationships among platelet responses: studies on the burst in proton liberation, lactate production, and oxygen uptake during platelet aggregation and Ca2+ secretion. Blood. 57:956–966. doi:10.1182/blood.v57.5.956.bloodjournal575956.

3. Anders, S., P.T. Pyl, and W. Huber. 2015. HTSeq--a Python framework to work with high-throughput sequencing data. Bioinformatics. 31:166–169. doi:10.1093/bioinformatics/btu638.

4. Antony, P.M.A., O. Boyd, C. Trefois, W. Ammerlaan, M. Ostaszewski, A.S. Baumuratov, L. Longhino, L. Antunes, W. Koopman, R. Balling, and N.J. Diederich. 2015. Platelet mitochondrial membrane potential in Parkinson’s disease. Ann. Clin. Transl. Neurol. 2:67–73. doi:10.1002/acn3.151.

5. Avila, C., R. Huang, M. Stevens, A. Aponte, D. Tripodi, K. Kim, and M. Sack. 2012. Platelet Mitochondrial Dysfunction is Evident in Type 2 Diabetes in Association with Modifications of Mitochondrial Anti-Oxidant Stress Proteins. Exp. Clin. Endocrinol. Diabetes. 120:248–251. doi:10.1055/s-0031-1285833.

6. Bergmeier, W., T. Rabie, A. Strehl, C.L. Piffath, M. Prostredna, D.D. Wagner, and B. Nieswandt. 2004. GPVI down-regulation in murine platelets through metalloproteinase-dependent shedding. Thromb. Haemost. 91:951–958. doi:10.1160/TH03-12-0795.

7. Brixner, V., M. Cardoso, E. Spaepen, and E. Seifried. 2024. Impact of shelf-life extension on platelet availability: Results from an inventory management modeling study. Transfus. Med. Hemother. 51:393–401. doi:10.1159/000537700.

8. Chen, S., Y. Su, and J. Wang. 2013. ROS-mediated platelet generation: a microenvironment-dependent manner for megakaryocyte proliferation, differentiation, and maturation. Cell Death Dis. 4:e722. doi:10.1038/cddis.2013.253.

9. Ding, Y., X. Gui, X. Chu, Y. Sun, S. Zhang, H. Tong, W. Ju, Y. Li, Z. Sun, M. Xu, Z. Li, R.K. Andrews, E.E. Gardiner, L. Zeng, K. Xu, and J. Qiao. 2023. MTH1 protects platelet mitochondria from oxidative damage and regulates platelet function and thrombosis. Nat. Commun. 14:4829. doi:10.1038/s41467-023-40600-7.

10. Dobin, A., C.A. Davis, F. Schlesinger, J. Drenkow, C. Zaleski, S. Jha, P. Batut, M. Chaisson, and T.R. Gingeras. 2013. STAR: ultrafast universal RNA-seq aligner. Bioinformatics. 29:15–21. doi:10.1093/bioinformatics/bts635.

11. Frankish, A., M. Diekhans, A.-M. Ferreira, R. Johnson, I. Jungreis, J. Loveland, J.M. Mudge, C. Sisu, J. Wright, J. Armstrong, I. Barnes, A. Berry, A. Bignell, S. Carbonell Sala, J. Chrast, F. Cunningham, T. Di Domenico, S. Donaldson, I.T. Fiddes, C. García Girón, J.M. Gonzalez, T. Grego, M. Hardy, T. Hourlier, T. Hunt, O.G. Izuogu, J. Lagarde, F.J. Martin, L. Martínez, S. Mohanan, P. Muir, F.C.P. Navarro, A. Parker, B. Pei, F. Pozo, M. Ruffier, B.M. Schmitt, E. Stapleton, M.-M. Suner, I. Sycheva, B. Uszczynska-Ratajczak, J. Xu, A. Yates, D. Zerbino, Y. Zhang, B. Aken, J.S. Choudhary, M. Gerstein, R. Guigó, T.J.P. Hubbard, M. Kellis, B. Paten, A. Reymond, M.L. Tress, and P. Flicek. 2019. GENCODE reference annotation for the human and mouse genomes. Nucleic Acids Res. 47:D766–D773. doi:10.1093/nar/gky955.

12. Fujii, T., A. Sakata, S. Nishimura, K. Eto, and S. Nagata. 2015. TMEM16F is required for phosphatidylserine exposure and microparticle release in activated mouse platelets. Proc. Natl. Acad. Sci. U. S. A. 112:12800–12805. doi:10.1073/pnas.1516594112.

13. Gardiner, E.E., D. Karunakaran, Y. Shen, J.F. Arthur, R.K. Andrews, and M.C. Berndt. 2007. Controlled shedding of platelet glycoprotein (GP)VI and GPIb-IX-V by ADAM family metalloproteinases. J. Thromb. Haemost. 5:1530–1537. doi:10.1111/j.1538-7836.2007.02590.x.

14. Hirata, S., T. Murata, D. Suzuki, S. Nakamura, R. Jono-Ohnishi, H. Hirose, A. Sawaguchi, S. Nishimura, N. Sugimoto, and K. Eto. 2017. Selective Inhibition of ADAM17 Efficiently Mediates Glycoprotein Ibα Retention During Ex Vivo Generation of Human Induced Pluripotent Stem Cell-Derived Platelets. Stem Cells Transl. Med. 6:720–730. doi:10.5966/sctm.2016-0104.

15. Ito, Y., S. Nakamura, N. Sugimoto, T. Shigemori, Y. Kato, M. Ohno, S. Sakuma, K. Ito, H. Kumon, H. Hirose, H. Okamoto, M. Nogawa, M. Iwasaki, S. Kihara, K. Fujio, T. Matsumoto, N. Higashi, K. Hashimoto, A. Sawaguchi, K.I. Harimoto, M. Nakagawa, T. Yamamoto, M. Handa, N. Watanabe, E. Nishi, F. Arai, S. Nishimura, and K. Eto. 2018. Turbulence Activates Platelet Biogenesis to Enable Clinical Scale Ex Vivo Production. Cell. 174:636–648.e18. doi:10.1016/j.cell.2018.06.011.

16. Jacob, S., Y. Kosaka, S. Bhatlekar, F. Denorme, H. Benzon, A. Moody, V. Moody, E.A. Tugolukova, G. Hull, N. Kishimoto, B.K. Manne, L. Guo, R. Souvenir, B.J. Seliger, A.S. Eustes, K. Hoerger, N.D. Tolley, A.N. Fatahian, S. Boudina, D.C. Christiani, Y. Wei, C. Ju, R.A. Campbell, M.T. Rondina, E.D. Abel, P.F. Bray, A.S. Weyrich, and J.W. Rowley. 2024. Mitofusin-2 regulates platelet mitochondria and function. Circ. Res. 134:143–161. doi:10.1161/CIRCRESAHA.123.322914.

17. Jansen, A.J.G., E.C. Josefsson, V. Rumjantseva, Q.P. Liu, H. Falet, W. Bergmeier, S.M. Cifuni, R. Sackstein, U.H. von Andrian, D.D. Wagner, J.H. Hartwig, and K.M. Hoffmeister. 2012. Desialylation accelerates platelet clearance after refrigeration and initiates GPIbα metalloproteinase-mediated cleavage in mice. Blood. 119:1263–1273. doi:10.1182/blood-2011-05-355628.

18. Jedlička, J., R. Kunc, and J. Kuncová. 2021. Mitochondrial respiration of human platelets in young adult and advanced age - Seahorse or O2k? Physiol. Res. 70:S369–S379. doi:10.33549/physiolres.934812.

19. Karasawa, S., T. Araki, M. Yamamoto-Hino, and A. Miyawaki. 2003. A green-emitting fluorescent protein from Galaxeidae coral and its monomeric version for use in fluorescent labeling. J. Biol. Chem. 278:34167–34171. doi:10.1074/jbc.M304063200.

20. Kechin, A., U. Boyarskikh, A. Kel, and M. Filipenko. 2017. CutPrimers: A new tool for accurate cutting of primers from reads of targeted next generation sequencing. J. Comput. Biol. 24:1138–1143. doi:10.1089/cmb.2017.0096.

21. Kholmukhamedov, A., R. Janecke, H.-J. Choo, and S.M. Jobe. 2018. The mitochondrial calcium uniporter regulates procoagulant platelet formation. J. Thromb. Haemost. 16:2315–2321. doi:10.1111/jth.14284.

22. Lefrançais, E., G. Ortiz-Muñoz, A. Caudrillier, B. Mallavia, F. Liu, D.M. Sayah, E.E. Thornton, M.B. Headley, T. David, S.R. Coughlin, M.F. Krummel, A.D. Leavitt, E. Passegué, and M.R. Looney. 2017. The lung is a site of platelet biogenesis and a reservoir for haematopoietic progenitors. Nature. 544:105–109. doi:10.1038/nature21706.

23. Ludeman, M.J., Y.W. Zheng, K. Ishii, and S.R. Coughlin. 2004. Regulated shedding of PAR1 N-terminal exodomain from endothelial cells. J. Biol. Chem. 279:18592–18599. doi:10.1074/jbc.M310836200.

24. Melchinger, H., K. Jain, T. Tyagi, and J. Hwa. 2019. Role of Platelet Mitochondria: Life in a Nucleus-Free Zone. Frontiers in Cardiovascular Medicine. 6:1–11. doi:10.3389/fcvm.2019.00153.

25. Merino, F., S. Pospich, and S. Raunser. 2020. Towards a structural understanding of the remodeling of the actin cytoskeleton. Semin. Cell Dev. Biol. 102:51–64. doi:10.1016/j.semcdb.2019.11.018.

26. Metz, V.V., E. Kojro, D. Rat, and R. Postina. 2012. Induction of RAGE shedding by activation of G protein-coupled receptors. PLoS One. 7:e41823. doi:10.1371/journal.pone.0041823.

27. Milroy, L.-G., S. Rizzo, A. Calderon, B. Ellinger, S. Erdmann, J. Mondry, P. Verveer, P. Bastiaens, H. Waldmann, L. Dehmelt, and H.-D. Arndt. 2012. Selective chemical imaging of static actin in live cells. J. Am. Chem. Soc. 134:8480–8486. doi:10.1021/ja211708z.

28. Miyata, Y., and K. Segawa. 2022. Protocol to analyze lipid asymmetry in the plasma membrane. STAR Protoc. 3:101870. doi:10.1016/j.xpro.2022.101870.

29. Moore, A.S., S.M. Coscia, C.L. Simpson, F.E. Ortega, E.C. Wait, J.M. Heddleston, J.J. Nirschl, C.J. Obara, P. Guedes-Dias, C.A. Boecker, T.L. Chew, J.A. Theriot, J. Lippincott-Schwartz, and E.L.F. Holzbaur. 2021. Actin cables and comet tails organize mitochondrial networks in mitosis. Nature. 591:659–664. doi:10.1038/s41586-021-03309-5.

30. Moreau, T., A.L. Evans, L. Vasquez, M.R. Tijssen, Y. Yan, M.W. Trotter, D. Howard, M. Colzani, M. Arumugam, W.H. Wu, A. Dalby, R. Lampela, G. Bouet, C.M. Hobbs, D.C. Pask, H. Payne, T. Ponomaryov, A. Brill, N. Soranzo, W.H. Ouwehand, R.A. Pedersen, and C. Ghevaert. 2016. Large-scale production of megakaryocytes from human pluripotent stem cells by chemically defined forward programming. Nat. Commun. 7. doi:10.1038/ncomms11208.

31. Nakamura, S., N. Takayama, S. Hirata, H. Seo, H. Endo, K. Ochi, K.I. Fujita, T. Koike, K.I. Harimoto, T. Dohda, A. Watanabe, K. Okita, N. Takahashi, A. Sawaguchi, S. Yamanaka, H. Nakauchi, S. Nishimura, and K. Eto. 2014. Expandable megakaryocyte cell lines enable clinically applicable generation of platelets from human induced pluripotent stem cells. Cell Stem Cell. 14:535–548. doi:10.1016/j.stem.2014.01.011.

32. Nishikii, H., K. Eto, N. Tamura, K. Hattori, B. Heissig, T. Kanaji, A. Sawaguchi, S. Goto, J. Ware, and H. Nakauchi. 2008. Metalloproteinase regulation improves in vitro generation of efficacious platelets from mouse embryonic stem cells. J. Exp. Med. 205:1917–1927. doi:10.1084/jem.20071482.

33. Nishimura, S., M. Nagasaki, S. Kunishima, A. Sawaguchi, A. Sakata, H. Sakaguchi, T. Ohmori, I. Manabe, J.E. Italiano, T. Ryu, N. Takayama, I. Komuro, T. Kadowaki, K. Eto, and R. Nagai. 2015. IL-1α induces thrombopoiesis through megakaryocyte rupture in response to acute platelet needs. J. Cell Biol. 209:453–466. doi:10.1083/jcb.201410052.

34. Okamoto, H., K. Fujio, S. Nakamura, Y. Harada, H. Hayashi, N. Higashi, A. Ninomiya, R. Tanaka, N. Sugimoto, N. Takayama, A. Kaneda, A. Sawaguchi, Y. Kato, and K. Eto. 2024. Defective flow space limits the scaling up of turbulence bioreactors for platelet generation. Communications Engineering. 3:1–12. doi:10.1038/s44172-024-00219-y.

35. Panch, S.R., L. Guo, and R. Vassallo. 2023. Platelet transfusion refractoriness due to HLA alloimmunization: Evolving paradigms in mechanisms and management. Blood Rev. 62:101135. doi:10.1016/j.blre.2023.101135.

36. Pospich, S., F. Merino, and S. Raunser. 2020. Structural effects and functional implications of phalloidin and jasplakinolide binding to actin filaments. Structure. 28:437–449.e5. doi:10.1016/j.str.2020.01.014.

37. Potts, K.S., A. Farley, C.A. Dawson, J. Rimes, C. Biben, C.D. Graaf, M.A. Potts, O.J. Stonehouse, A. Carmagnac, P. Gangatirkar, E.C. Josefsson, C. Anttila, D. Amann-Zalcenstein, S. Naik, W.S. Alexander, D.J. Hilton, E.D. Hawkins, and S. Taoudi. 2020. Membrane budding is a major mechanism of in vivo platelet biogenesis. J. Exp. Med. 217. doi:10.1084/JEM.20191206.

38. Richardson, J.L., R.A. Shivdasani, C. Boers, J.H. Hartwig, and J.E. Italiano. 2005. Mechanisms of organelle transport and capture along proplatelets during platelet production. Blood. 106:4066–4075. doi:10.1182/blood-2005-06-2206.

39. Richman, T.R., J.A. Ermer, J. Baker, S.J. Siira, B.T. Kile, M.D. Linden, O. Rackham, and A. Filipovska. 2023. Mitochondrial gene expression is required for platelet function and blood clotting. Cell Rep. 42:113312. doi:10.1016/j.celrep.2023.113312.

40. Shi, C., K. Guo, D.T. Yew, Z. Yao, E.L. Forster, H. Wang, and J. Xu. 2008. Effects of ageing and Alzheimer’s disease on mitochondrial function of human platelets. Exp. Gerontol. 43:589–594. doi:10.1016/j.exger.2008.02.004.

41. Sim, X., D. Jarocha, V. Hayes, H.A. Hanby, M.S. Marks, R.M. Camire, D.L. French, M. Poncz, and P. Gadue. 2017. Identifying and enriching platelet-producing human Stem cell-derived megakaryocytes using factor V uptake. Blood. 130:192–204. doi:10.1182/blood-2017-01-761049.

42. Sommer, A., F. Kordowski, J. Büch, T. Maretzky, A. Evers, J. Andrä, S. Düsterhöft, M. Michalek, I. Lorenzen, P. Somasundaram, A. Tholey, F.D. Sönnichsen, K. Kunzelmann, L. Heinbockel, C. Nehls, T. Gutsmann, J. Grötzinger, S. Bhakdi, and K. Reiss. 2016. Phosphatidylserine exposure is required for ADAM17 sheddase function. Nat. Commun. 7:11523. doi:10.1038/ncomms11523.

43. Sone, M., S. Nakamura, S. Umeda, H. Ginya, M. Oshima, M.A. Kanashiro, S.K. Paul, K. Hashimoto, E. Nakamura, Y. Harada, K. Tsujimura, A. Saraya, T. Yamaguchi, N. Sugimoto, A. Sawaguchi, A. Iwama, K. Eto, and N. Takayama. 2021. Silencing of p53 and CDKN1A establishes sustainable immortalized megakaryocyte progenitor cells from human iPSCs. Stem Cell Reports. 16:2861–2870. doi:10.1016/j.stemcr.2021.11.001.

44. Sugimoto, N., J. Kanda, S. Nakamura, T. Kitano, M. Hishizawa, T. Kondo, S. Shimizu, A. Shigemasa, H. Hirai, Y. Arai, M. Minami, H. Tada, D. Momose, K.-R. Koh, M. Nogawa, N. Watanabe, S. Okamoto, M. Handa, A. Sawaguchi, N. Matsuyama, M. Tanaka, T. Hayashi, A. Fuchizaki, Y. Tani, A. Takaori-Kondo, and K. Eto. 2022a. iPLAT1: the first-in-human clinical trial of iPSC-derived platelets as a phase 1 autologous transfusion study. Blood. 140:2398–2402. doi:10.1182/blood.2022017296.

45. Sugimoto, N., S. Nakamura, S. Shimizu, A. Shigemasa, J. Kanda, N. Matsuyama, M. Tanaka, T. Hayashi, A. Fuchizaki, M. Nogawa, N. Watanabe, S. Okamoto, M. Handa, A. Sawaguchi, D. Momose, K.-R. Koh, Y. Tani, A. Takaori-Kondo, and K. Eto. 2022b. Production and nonclinical evaluation of an autologous iPSC-derived platelet product for the iPLAT1 clinical trial. Blood Adv. 6:6056–6069. doi:10.1182/bloodadvances.2022008512.

46. Tilburg, J., I.C. Becker, and J.E. Italiano. 2022. Don’t you forget about me(gakaryocytes). Blood. 139:3245–3254. doi:10.1182/blood.2020009302.

47. Twomey, L.C., R.G. Wallace, P.M. Cummins, B. Degryse, S. Sheridan, M. Harrison, N. Moyna, G. Meade-Murphy, N. Navasiolava, M. Custaud, and R.P. Murphy. 2018. Platelets: From formation to function. Homeost. Health Dis. doi:10.5772/INTECHOPEN.80924.

48. Wang, L., Q. Wu, Z. Fan, R. Xie, Z. Wang, and Y. Lu. 2017. Platelet mitochondrial dysfunction and the correlation with human diseases. Biochem. Soc. Trans. 45:1213–1223. doi:10.1042/BST20170291.

49. Zhao, X., D. Alibhai, T.G. Walsh, N. Tarassova, M. Englert, S.Z. Birol, Y. Li, C.M. Williams, C.R. Neal, P. Burkard, S.J. Cross, E.W. Aitken, A.K. Waller, J.B. Beltrán, P.W. Gunning, E.C. Hardeman, E.O. Agbani, B. Nieswandt, I. Hers, C. Ghevaert, and A.W. Poole. 2023. Highly efficient platelet generation in lung vasculature reproduced by microfluidics. Nat. Commun. 14:4026. doi:10.1038/s41467-023-39598-9.

50. Zharikov, S., and S. Shiva. 2013. Platelet mitochondrial function: From regulation of thrombosis to biomarker of disease. Biochem. Soc. Trans. 41:118–123. doi:10.1042/BST20120327.

51. Zhou, K., Y. Xia, M. Yang, W. Xiao, L. Zhao, R. Hu, K.M. Shoaib, R. Yan, and K. Dai. 2022. Actin polymerization regulates glycoprotein Ibα shedding. Platelets. 33:381–389. doi:10.1080/09537104.2021.1922882.

