## Supplementary figure legends for "Turbulence orchestrates actin-mitochondria dynamics to preserve GPIbα and support iPSC-derived platelet biogenesis"

**Supplemental Figure 1. iPSC-derived platelets (iPSC-PLTs) produced under turbulent conditions exhibit superior quality compared to those from static cultures (related to Figure 1)**

(A) Schematic diagram of doxycycline (DOX)-ON expansion and DOX-OFF maturation culture phases. All data were collected from day 0 to day 6 of the DOX-OFF phase.

(B) Representative flow cytometry plots showing P-selectin expression and PAC-1 binding (left) and bar graphs quantifying the percentage of PAC-1$^+$/P-selectin$^+$ (CD62P$^+$) iPSC-PLTs, either unstimulated (black) or stimulated (grey) with ADP/TRAP6 (top) or PMA (bottom) in dish, flask, or VerMES cultures (n = 3; error bars represent SEM; unpaired two-tailed t-test: ns, not significant; $**p < 0.01$; $***p < 0.001$).

(C) Bar graph showing annexin V binding in day 6 iPSC-PLTs cultured in dish, flask, or VerMES conditions (n = 3; mean ± SEM; unpaired two-tailed t-test: ns; $**p < 0.01$; $***p < 0.001$).

(D) Representative flow cytometry plots showing GPIbα and PAR-1 or GPVI expression in day 6 iPSC-PLTs from dish, flask, or VerMES cultures.

(E) Representative flow cytometry plots showing CD41 and GPIbα expression in iPSC-PLTs from day 1 to day 6 of culture in dish, flask, or VerMES conditions. The gate highlights GPIbα$^+$ (top, red) and GPIbα$^-$ (bottom, black) populations.

(F) Quantification of GPIbα$^+$ platelet numbers from day 1 to day 6 of imMKCL maturation in dish (orange), flask (purple), or VerMES (green) cultures. Data are shown as mean ± SEM (n = 3).

(G–I) Effects of KP457 supplementation on platelet yield, annexin V binding, and activation marker expression in day 6 iPSC-PLTs.

(G) Platelet yield per imMKCL in dish or flask cultures with or without KP457 treatment.

(H) Annexin V binding in the presence (blue) or absence (red) of KP457.

(I) P-selectin (CD62P) expression and PAC-1 binding after ADP/TRAP6 stimulation in platelets cultured with or without KP457.

(J–L) Factor V uptake by maturing imMKCLs and its transfer to iPSC-PLTs in dish cultures.

(J) Schematic showing the addition of Alexa Fluor 488–labeled factor V to maturing imMKCLs and its transfer to iPSC-PLTs.

(K) Representative flow cytometry plots showing GPIbα expression (left) and factor V fluorescence (right) in day 4 imMKCLs.

(L) Representative histograms showing factor V fluorescence in day 6 iPSC-PLTs.

(M) Representative flow cytometry plots showing factor V$^+$ (red) and factor V$^-$ (blue) iPSC-PLTs analyzed for GPIbα, PAR-1, GPVI, mitochondrial membrane potential (MitoTracker Deep Red or MT-1), reactive oxygen species (CellROX), and ATP (BioTracker ATP Red).

**Supplemental Figure 2. Evaluation of mitochondrial functions and quality in iPSC-derived and stored human platelets**

(A) Vector map of the construct used in Figure 2C, with a mitochondria-targeted mAzami Green (CoralHue® Mito-mAG) expression cassette.

(B) Confocal micrographs of day 6 iPSC-derived platelets cultured in a dish or flask, plated on poly-L-lysine–coated Ibidi μ-dishes (quad format), and stained for CD41 (red), GPIbα (yellow),

and mitochondria (green). Images were acquired using a Carl Zeiss LSM900 confocal microscope with Airyscan. Scale bar, 10 µm.
(C) Transmission electron microscopy (TEM) images showing mitochondrial density in day 6 iPSC-derived platelets cultured in a dish (top) or flask (bottom). Mitochondria are indicated by red spots.
(D–G) Assessment of mitochondrial function and viability in stored human platelets.
(D) Schematic diagram outlining the experimental timeline. Human platelets were stored at 22 °C or 37 °C from day 1 to day 12; assays were performed between days 5 and 12.
(E) Representative flow cytometry plots and histograms showing GPIbα expression, mitochondrial membrane potential (MitoTracker Deep Red), and annexin V binding in human platelets on days 5, 8, and 12 following storage at either 22 °C or 37 °C.
(F) Representative histograms showing GPIbα expression and mitochondrial membrane potential (MitoTracker Deep Red) before (gray) and after FCCP treatment, with (blue) or without (red) KP457 supplementation.
(G) Representative flow cytometry plots showing GPIbα and GPVI expression, along with mitochondrial membrane potential (MitoTracker Deep Red), in day 12 human platelets treated with DMSO or supplemented with KP457, GI254023, TAPI-1, or GM6001.

**Supplemental Figure 3. Analysis of phospholipid asymmetry in iPSC-derived and human platelets**
(A) Representative flow cytometry plots showing surface GPIbα expression and fluorescence of either NBD–phosphatidylserine (NBD-PS, top) or NBD–phosphatidylcholine (NBD-PC, bottom) in day 6 iPSC-derived platelets cultured in a flask (left) or dish (right). Conditions include no BSA wash ($BSA^{-}$), after BSA wash ($BSA^{+}$), and after ionomycin treatment.
(B) Representative flow cytometry plots showing uptake of NBD-PS and NBD-PC by human platelets on days 6, 8, and 11.

**Supplemental Figure 4. Temporal analysis of surface marker expression and mitochondrial membrane potential in maturing imMKCLs**
(A, B) Flow cytometric analysis of CD41, GPIbα, and mitochondrial membrane potential (MitoTracker Deep Red) in imMKCLs cultured in a dish, flask, or VerMES from day 1 to day 6.
(A) Representative flow cytometry plots showing surface expression of CD41 and GPIbα over time.
(B) Representative plots showing changes in mitochondrial membrane potential during culture.
