## Supplementary figures for "Turbulence orchestrates actin-mitochondria dynamics to preserve GPIbα and support iPSC-derived platelet biogenesis"

### Supplementary Figure 1

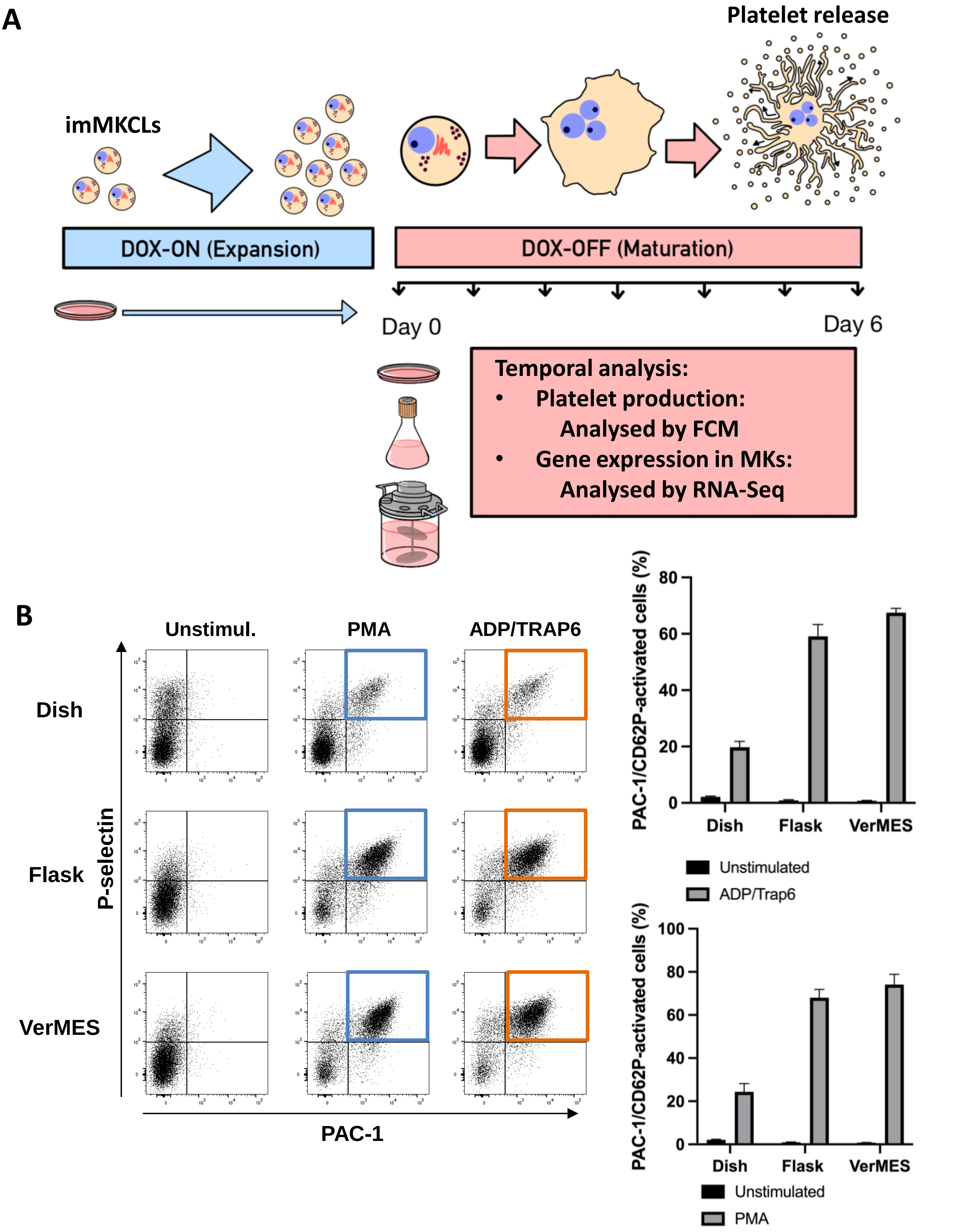

### Supplementary Figure 1

C

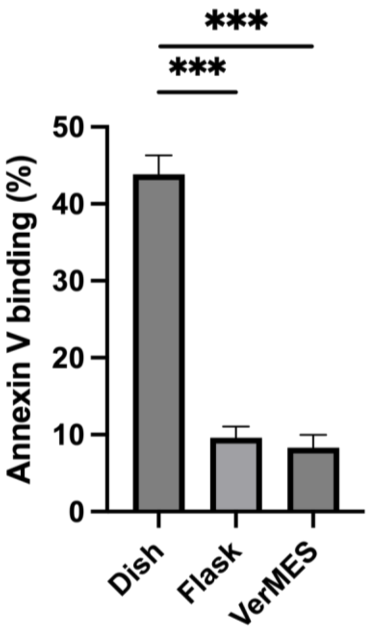


D

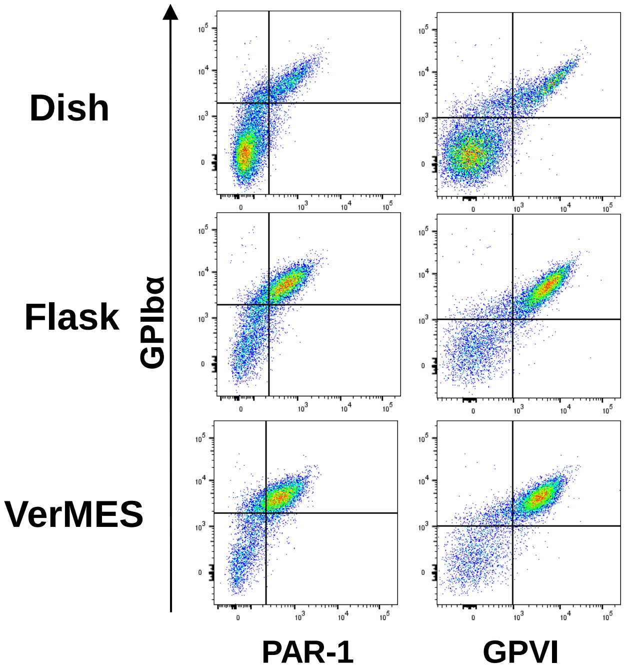


E

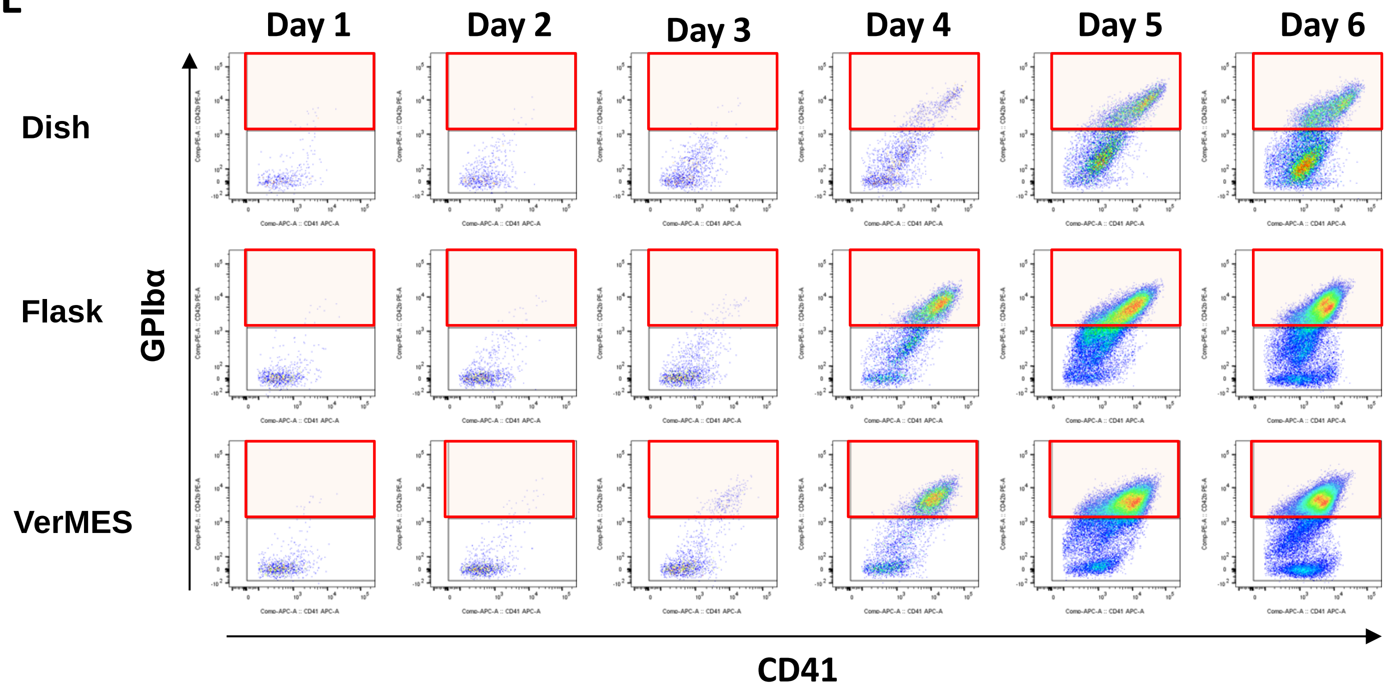

### Supplementary Figure 1

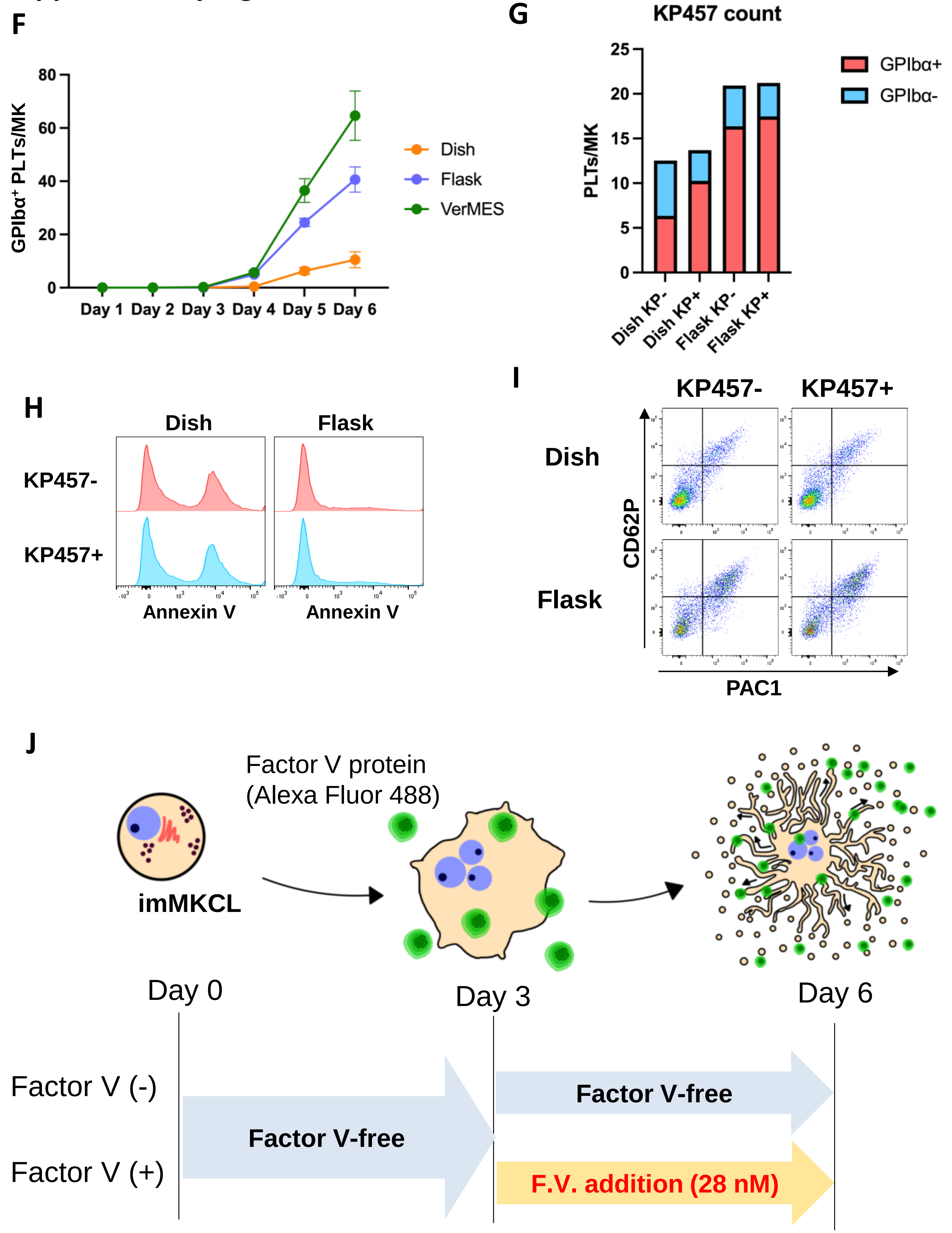

### Supplementary Figure 1

K

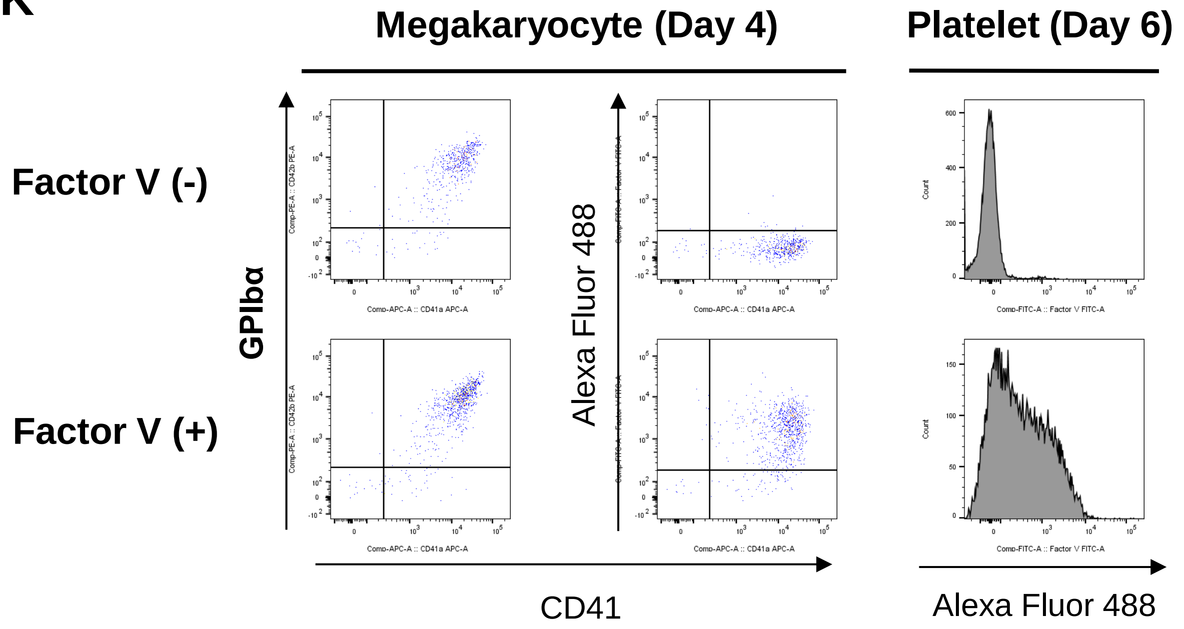


L

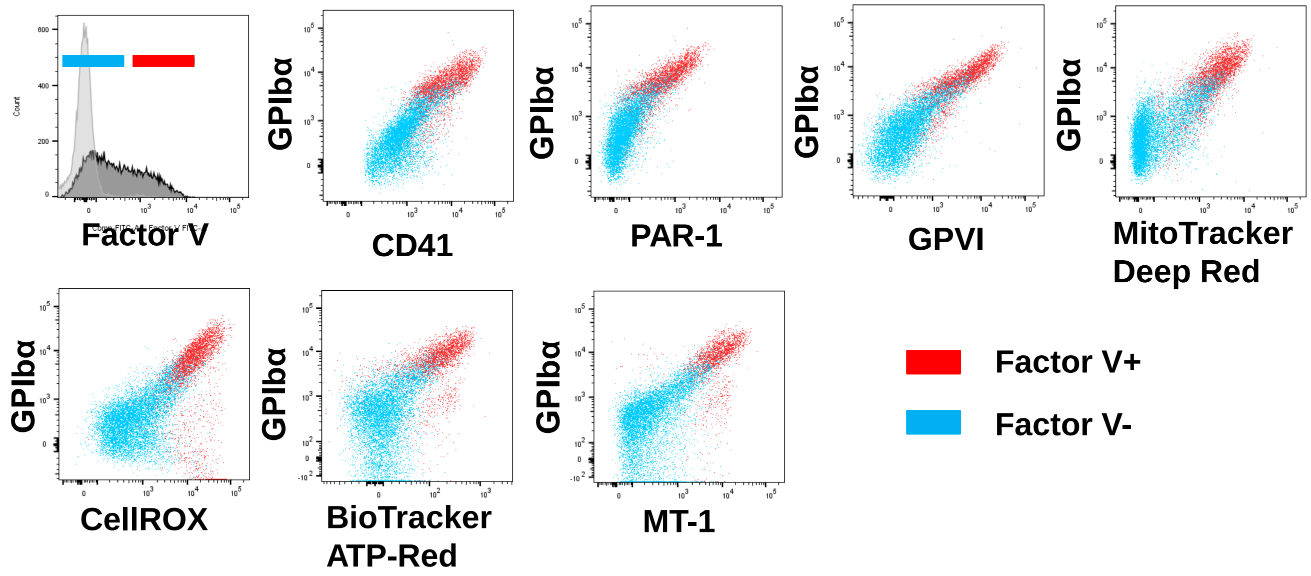

### Supplementary Figure 2

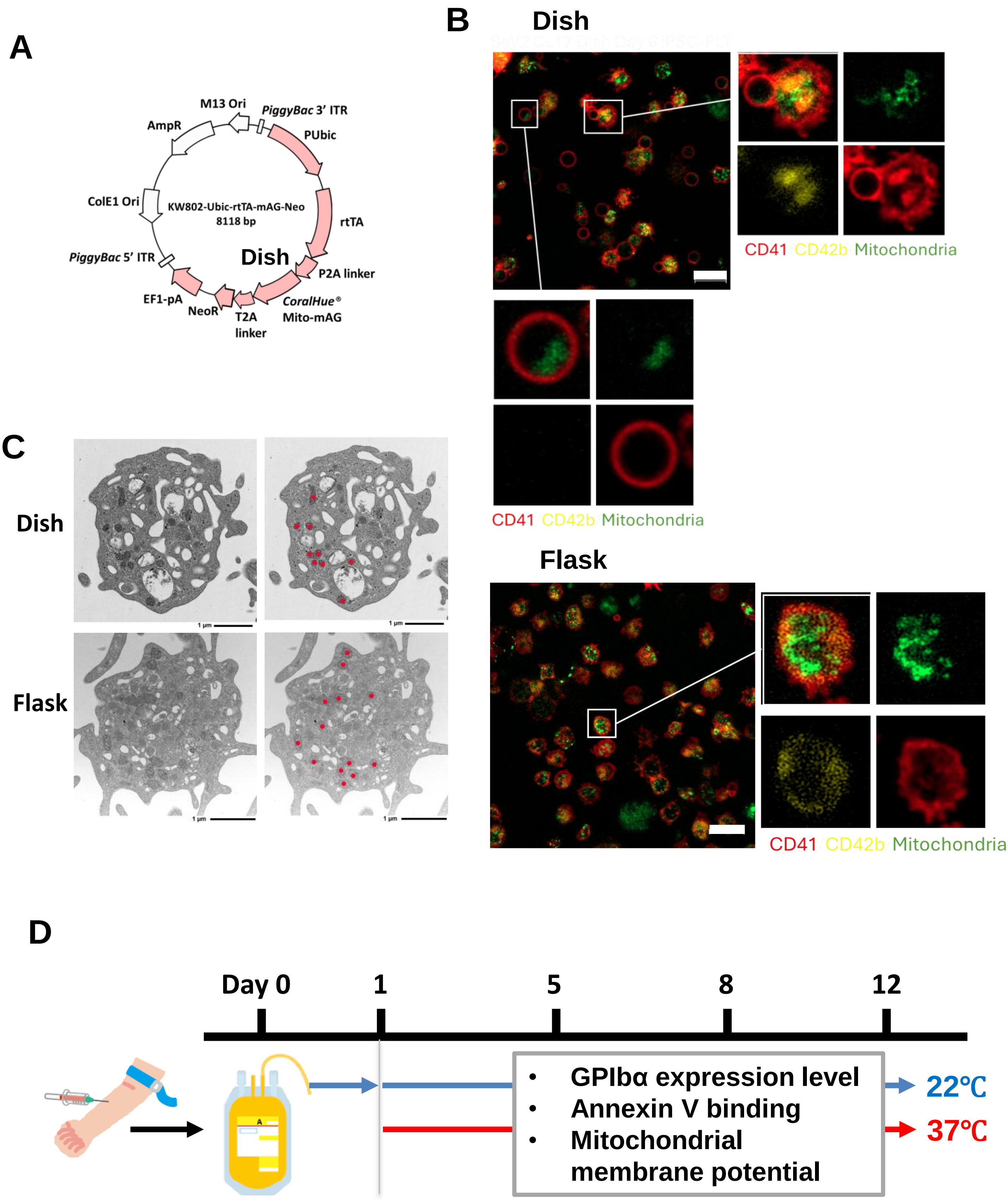

### Supplementary Figure 2

E

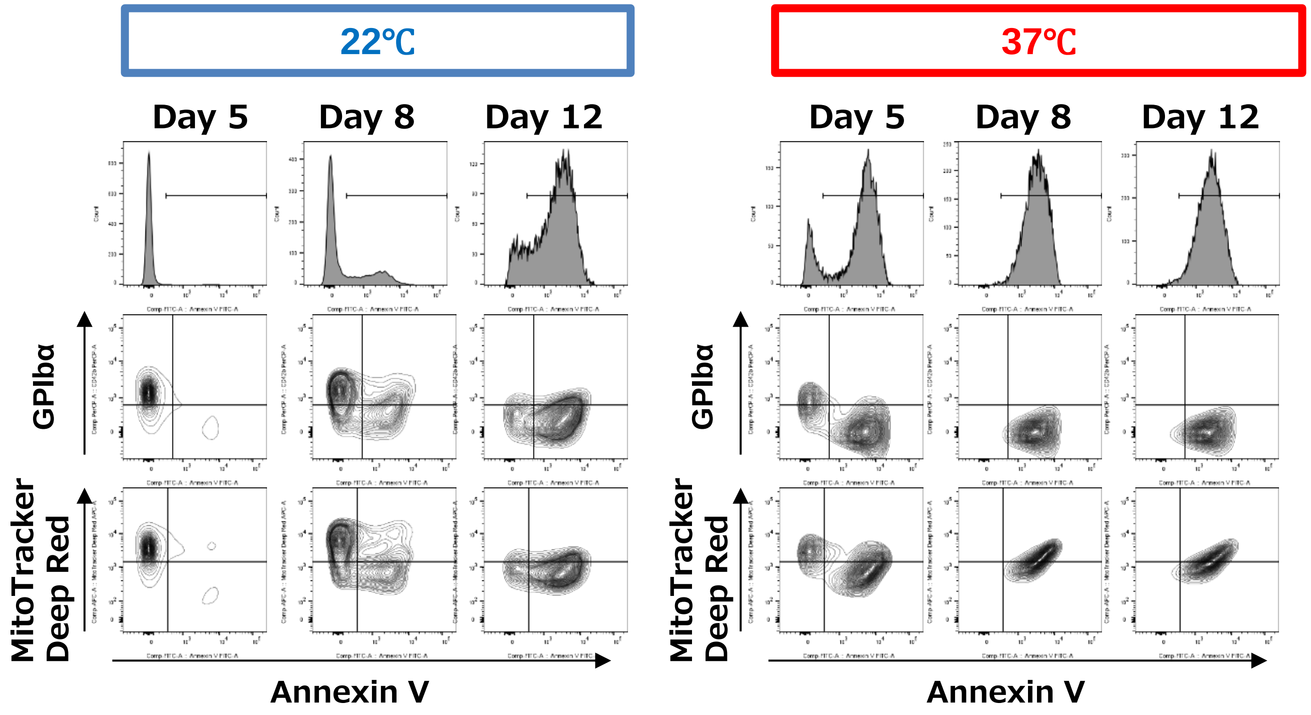


F

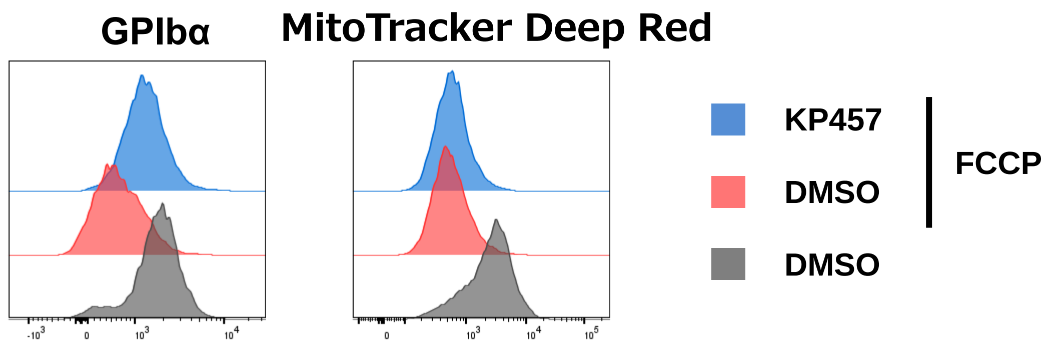


G

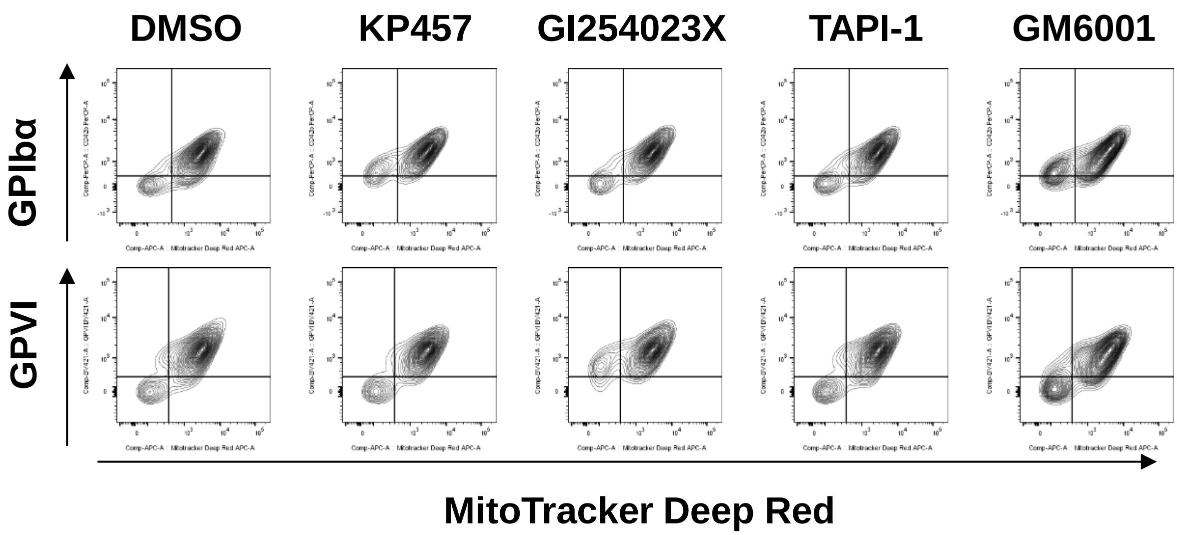

### Supplementary Figure 3

**A**

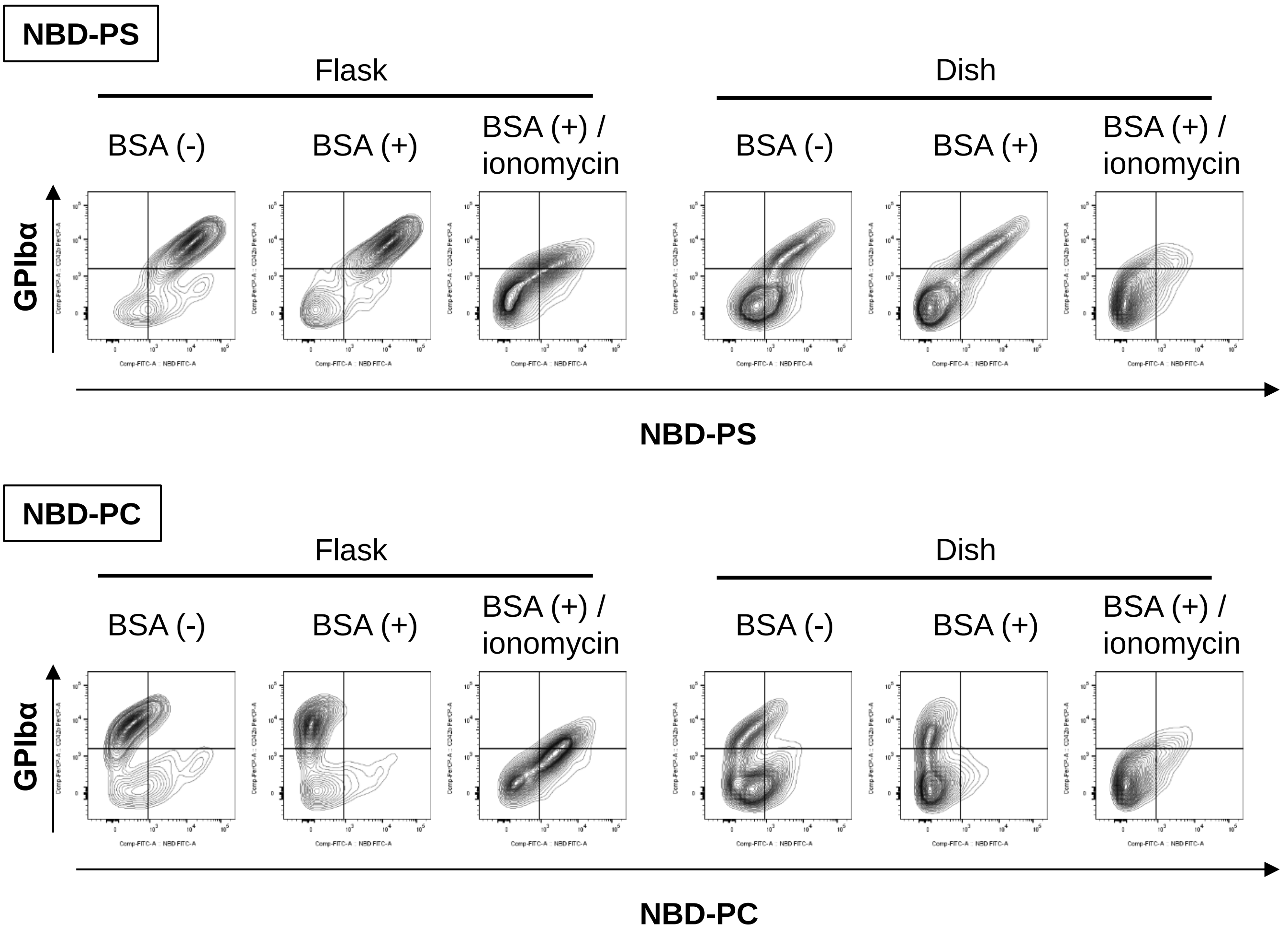


**B** **Human Platelet**

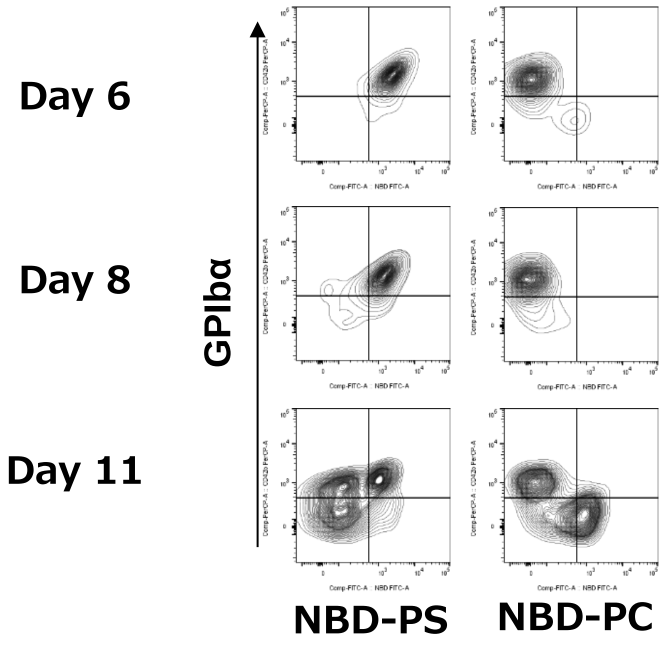

### Supplementary Figure 4

#### A Surface marker expression

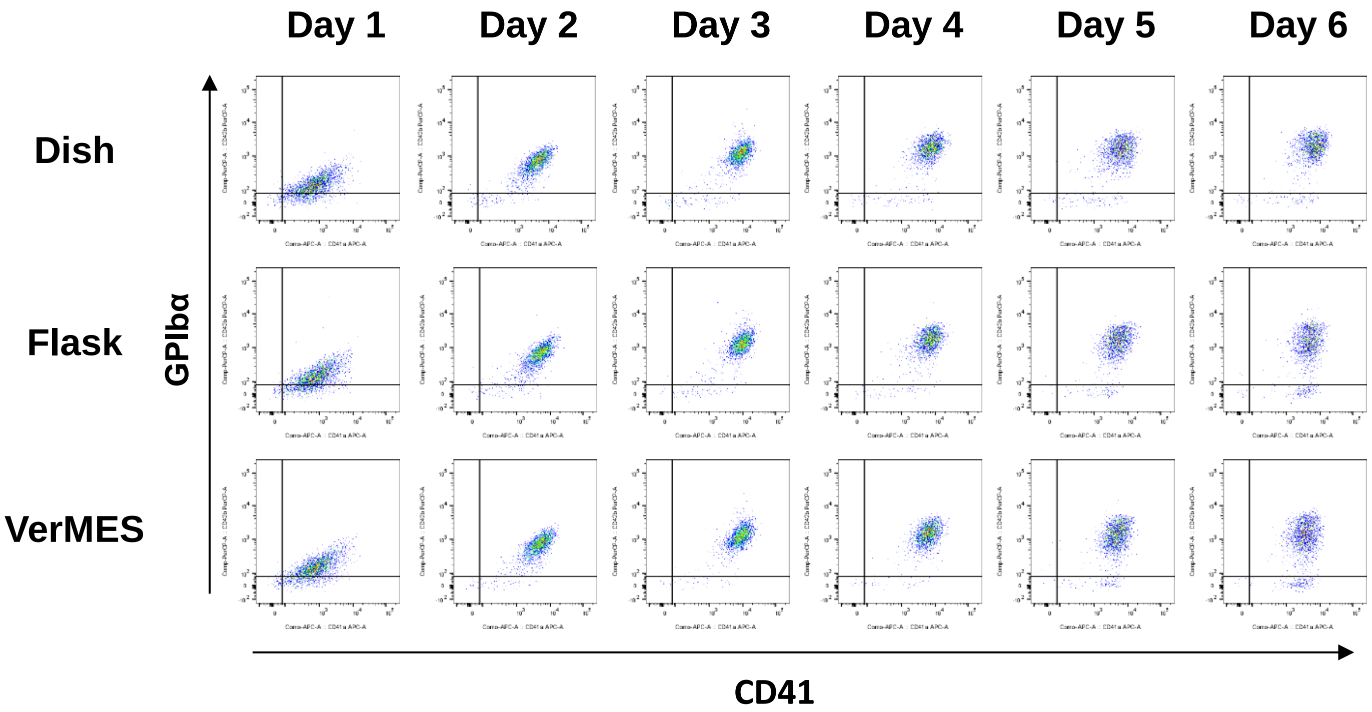


#### B Mitochondrial membrane potential

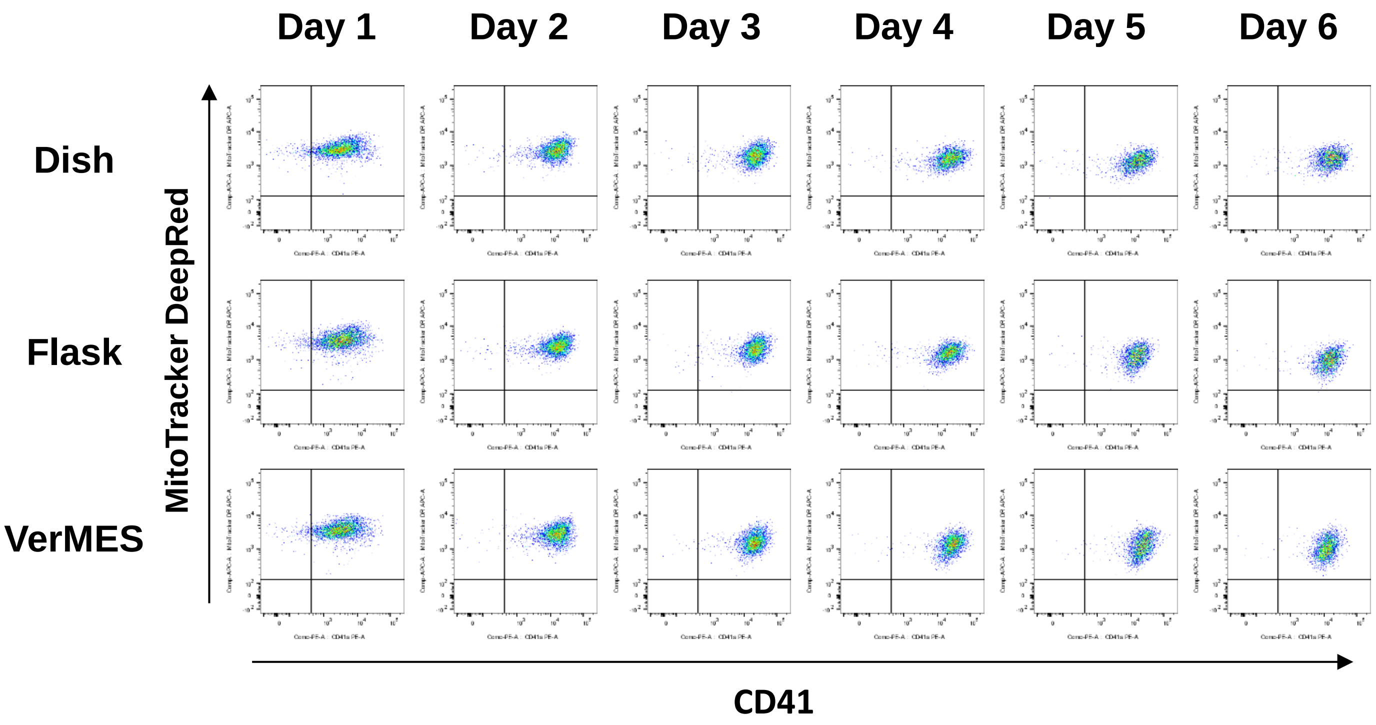
